# Fluorescent peptides for membrane tension and domain structure reporting

**DOI:** 10.64898/2025.12.30.697119

**Authors:** Thomas Suchyna, Jake Rossetto, Hannah Cetuk, Andriy Anishkin, Sergei Sukharev

## Abstract

Cell mechanics play significant roles in all aspects of cell function. While many types of fluorescent linear force sensors inserted in cellular fibrillar elements have been developed, few tools are available to track two-dimensional tension in cell membranes. Here, we present a novel principle for tension detection using a fluorescent probe based on the scaffold of the GsMTx4 peptide from *Grammostola* venom. Previously, we have shown that amphipathic GsMTx4 binds to lipids and inhibits mechanosensitive channels by inserting more deeply into the membrane at tensions near activation thresholds, thereby acting as a buffer clamping lateral pressure in the bilayer. We leverage this property of GsMTx4 to redistribute between the ‘shallow’ and ‘deep’ immersion states, thereby designing probes with a fluorescent moiety that increases quantum yield in nonpolar environments. GsMTx4 analogs carrying fluorescent groups at the two positions increase fluorescence intensity in osmotically shocked liposomes and aspirated giant vesicles in a near-linear fashion in response to physiological bilayer tensions. The responses show dependence on membrane composition, particularly lipid charge and the presence of lipid-ordering components, such as sphingomyelin and cholesterol. Langmuir compression isotherms recorded in the presence of NBD analogs indicated initial incorporation into the monolayer, followed by sharp expulsion at the monolayer-bilayer equivalence pressure, with correlated changes in monolayer compressibility and fluorescence, illustrating the basic principle of probe action. The probes show promise for monitoring tension in biological membranes at low, non-inhibitory concentrations. Experiments with native cell-derived membrane vesicles reveal heterogeneous baseline staining and tension responses, underscoring the probes’ selectivity for distinct membrane domains.

**Significance:** Cell mechanics are crucial for all cell functions, including division, survival, migration, and differentiation. Although many versions of fluorescent linear force sensors have been developed for cytoskeletal and ECM elements, few tools exist to monitor two-dimensional tension in cell membranes. Many cells are motile, actively deforming their membrane, supported and driven by the underlying cytoskeleton. There is a two-order-of-magnitude discrepancy between membrane tension estimates from the tether formation technique and the tensions that activate common mechanosensitive channels in most cells. This discrepancy highlights the need for non-invasive membrane probes that can independently measure membrane tension, especially since it can be highly localized and dynamic. Here, we introduce such probes and a new principle for tension measurement.

---

The generation of forces and the active movement are integral to every cellular process, including basic progression through the cell cycle, cell migration, adhesion to a substrate, and contractility. Mechanobiology is gaining prominence in all facets of normal growth and development, tissue repair, synapse formation between neurons or immune cells, pathogen invasion, cancer growth, and metastasis. The ability to detect and measure in situ forces directly within cells and tissues supports many mechanistic inferences and conclusions. Several techniques have been developed to measure linear stresses in fibrillar components using fluorescent reporters inserted into cytoskeletal or extracellular matrix (ECM) elements (1–6). Despite progress in detecting and quantifying linear forces, the distribution, magnitudes, and dynamics of two-dimensional stresses, i.e., tension, in cell membranes remain elusive.

While bacterial, fungal, and plant cells contain internal turgor pressure by building external cell walls (7–9), the mechanical integrity of wall-less animal cells relies on a cross-linked internal cytoskeleton that lines the membrane from within. This scaffolding is crucial; a simple example shows that an ‘unarmored’ membrane vesicle 1 µm in diameter, holding a modest osmotic gradient of 20 mOsm, experiences a nearly lytic tension of 12 mN/m. Osmotic gradients across the plasma membrane of an animal cell can be much greater, so the membranes must be mechanically shielded. To prevent high tensile stress, animal cells typically produce a significant excess of membrane, which is stored in protrusions, invaginations (10), and especially in caveolae (11), allowing moderate swelling to unfold the membrane without generating global tension. The question remains whether, without complete unfolding to its limits, the membrane ever experiences significant local tension. Evidently, it does, as the cells are exposed to external mechanical stresses and actively perturb their surface by moving, contracting, and pulling on cytoskeletal attachment points.

Estimates of membrane tension in mammalian cells based on tether pulling methods (12) range from 0.003 mN/m in neuronal growth cones (13) to 0.2-0.5 mN/m in rapidly moving keratinocytes (14). The equilibrium tether pulling force thus depends on the cell type, the state of the cytoskeleton, and motility. In many such experiments, the tether is pulled from a location outside the zone of active membrane or cytoskeletal dynamics (such as the leading edge), and the question is whether this method reliably reports local tension at other locations. Recent studies have shown that tension propagation (diffusion) across the cytoskeleton-supported membrane is extremely short-range (15), and for this reason, the tether-pulling force can be uncorrelated with the local tension at 2-3 μm. Detaching the membrane from the cytoskeleton (15) or swelling the cell osmotically (16) creates stronger mechanical coupling between the probing tethers, although under unnatural conditions.

Is there an alternative way to predict the natural level of tension that cell membranes might experience? It is widely accepted that cells use mechanosensitive channels (MSCs) as natural tension sensors to detect local membrane stress. The widespread presence of tension-gated MSCs, not only in sensory neurons but also in most somatic cells, underscores the importance of tension as a key physiological parameter. With the exception of the auditory transduction channel (TMC), which is gated by protocadherin filamentous tip links (17), the common mammalian MSCs directly respond to tension in the lipid bilayer (18, 19). The most abundant are cationic (excitatory) Piezo1 and Piezo2 (20, 21) and hyperpolarizing (inhibitory) K2P channels (TREK 1 and 2, TRAAK) (22). Among these, Piezo1 performs the majority of mechanosensitive functions in the cardiovascular system, kidney, and bladder (20), whereas Piezo2 functions in sensory (DRG) neurons and peripheral afferent endings, including tactile receptors in the skin (23). The vivid functional phenotypes of global and tissue-specific knock-outs of Piezo channels (20, 24–26) attest to the importance of tension as a biological parameter in normal development and physiology.

The tension activation midpoints for excitatory Piezo1 range from 2 to 6 mN/m in various cellular contexts (27, 28). For inhibitory K^+^-selective K2P channels, including TREK1, TREK2, and TRAAK, the activation midpoints are reported to be between 4 and 6 mN/m (29). In single-cell experiments, Piezo channels are activated at the base of the pulled tether (15). Piezo1 channels are also active at the trailing edge of keratinocytes and regulate retraction in these rapidly moving cells (30). Therefore, the range of 0-6 mN/m—much higher than previously reported (10, 14, 31)—may represent typical physiological tensions that fluctuate across different locations and are detected by these channels. Do we have the means to measure these tensions independently using a non-invasive synthetic sensor?

*Laurdan,* a polarity-sensitive probe (32), and its derivatives (33) have been widely used to measure membrane lipid order. The spectral readout, quantified as a general fluorescence polarization (GP) parameter (34), reflects the polarity at the boundary between the aliphatic and headgroup regions of the lipid bilayer. Laurdan was used to report on membrane tension in osmotically shocked liposomes (35), and, indeed, its GP parameter decreased linearly by 5% across tensions from 0 to 3 mN/m (36). However, this parameter showed the opposite trend in osmotically swollen mammalian cells (37).

*Flipper probes* were developed for the optical characterization of lateral lipid packing (38), and have also been shown to report on membrane tension (39). The Flipper-TR dye has two dithienothiophene planar domains, and the averaged mutual orientation of these domains determines the fluorescence lifetime, varying from 3.6 ns in disordered bilayers (POPC) to 6.5 ns in more ordered sphingomyelin-cholesterol bilayers. According to (38), membrane tension causes lateral lipid chain compression and an increase in fluorescence lifetime. The dye sensitivity depends on membrane composition and reports on tensions up to 0.6 mN/m (38). The method requires advanced fluorescence lifetime imaging microscopy (FLIM).

Here, we propose a different principle for sensing membrane tension. Our tension probe is based on the peptide scaffold of the inhibitory GsMTx4 peptide, a known gating modifier of mechanosensitive channels, particularly Piezos (40, 41). The peptide was shown to modulate the gating of MS channels by adsorbing to the lipid membrane and transitioning from the ‘shallow’ to the ‘deep’ binding position under tension, thereby acting as a ‘mobile’ amphipathic component that inserts into the leaflet and buffers tension changes (42). The rigid peptide has a conical shape with a polar rim and a hydrophobic tip that contains two tryptophans, one of which has been replaced with the polarity-sensitive nitrobenzoxadiazole (NBD) fluorophore (43). In the resting membrane, the probe is in the aqueous phase or bound in a ‘shallow’ state with NBD positioned among polar headgroups. Under tension, the NBD probe penetrates more deeply into the nonpolar environment, resulting in a higher fluorescence quantum yield than in a polar environment. The rise in fluorescence indicates the onset of tension. Removing tension increases the lateral pressure on lipids, causing the probe to be expelled to its shallow position. The probe reports on a broad range of tensions while present in non-inhibitory concentrations.

We also show that probe partitioning and fluorescence changes depend on lipid context and can be used to characterize differential domains. Below, we describe the probe and test its properties in several model systems and native membranes.

## Computational probe design and simulations

The structure of the 35-amino acid GsMTx4 peptide, presented as the main chain and solvent-accessible surfaces (Fig.1), resembles an inverted cone with the downward-oriented tip formed by two tryptophan sidechains, W6 and W7 (44). These residues, along with L3, L26, and L29, and F5, F27, and F32, contribute to the hydrophobicity of the bottom face. The wider cone perimeter and the top face are polar and harbor multiple positive charges, including R18, K15, K20, K22, K25, and K28, which can interact with phospholipid phosphates. The two acidic residues, D13 and D14, on the upper face are expected to stabilize the tip-down orientation by repelling the phospholipid phosphates. The peptide shape is well-suited to wedge between the lipid headgroups and to immerse its tip into the polar/apolar boundary and further into the aliphatic core of the membrane (42, 45). The GsMTx4’s solvent-accessible surfaces (Figs. 1B and C) are colored by hydrophobicity and electrostatic potential. However, the acidic E4 and basic K9, located on the lower side, break the perfect separation of the hydrophobic and polar faces, suggesting that, in addition to the predicted deeply immersed conformation, there might be alternative membrane-bound states. Our simulations and experiments indicated that there are at least two modes of adsorption to a membrane (or lipid monolayer), deep and shallow, and lateral pressure changes drive the redistribution between these states (42).

**Figure 1.**
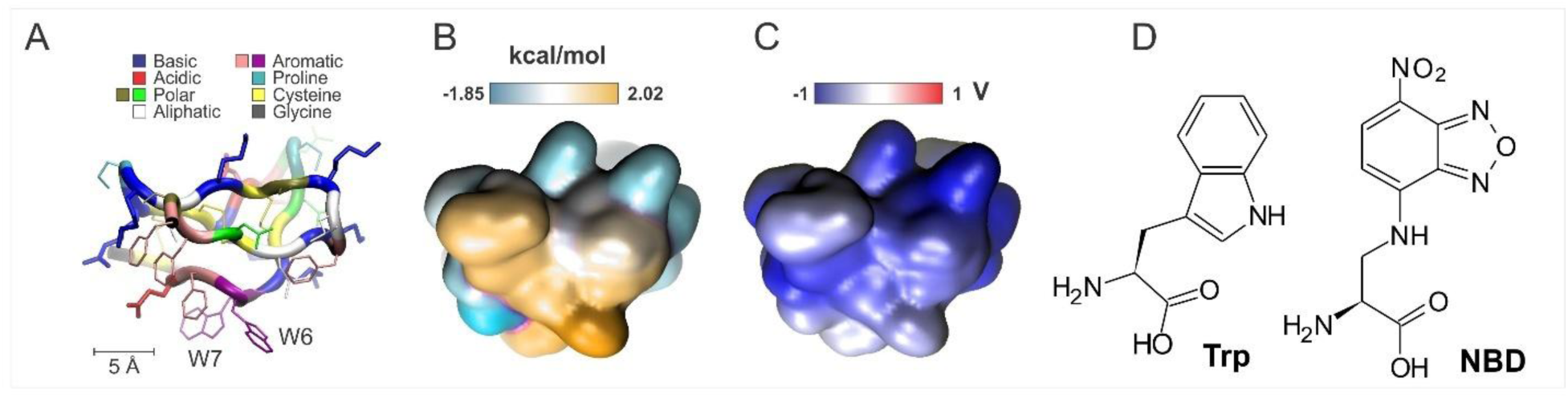
The structure of GsMTx4 and the design of the tension sensor. (A) The NMR structure of GsMTx4 (PDB 1TYK) is shown with critical side chains. The solvent-accessible surface of the peptide is colored by the hydrophobicity scale (B), characterizing affinity to the membrane interface (46), and by electrostatic potential(C). (D) Chemical structures of Tryptophan and NBD amino acid.

These considerations suggest that the tip of the cone would be the optimal location for a fluorophore sensitive to environmental polarity. GsMTx4 has two neighboring Trp residues at the hydrophobic face, W6 and W7. The endogenous tryptophan fluorescence, previously used to monitor membrane association and the depth of peptide penetration, exhibits a relatively weak resolving power (47, 48). To increase sensitivity, we chose NBD (7-nitrobenz-2-oxa-1,3-diazol-4-yl), one of the small, environment-sensitive dyes (463/536 nm), which resembles the side chain of tryptophan (Fig. 1D). NBD exhibits a ∼10 times higher quantum yield in an apolar environment, has a considerably longer lifetime, and displays moderate red-edge spectral shifts (REES) (43, 49). We did not use a cysteine or amino group-specific coupling to attach the fluorophore. Instead, we have incorporated a fluorescent amino acid with an NBD sidechain into two alternative positions using solid-phase peptide synthesis (by CS Bio Co.).

The results of tests presented below show that W6NBD is more sensitive to changes in lateral pressure/tension in extruded liposomes than W7NBD, but less sensitive in large vesicles. In this report, we present the majority of the data obtained with W6NBD, while in some experiments, we show W7NBD data for comparison.

To predict the behavior of the sensor peptides in lipid bilayers, we assembled membrane simulation systems comprising 128 lipids per leaflet and placed 4 W6NBD peptides per simulation cell (Supplemental Figure S1). Initial simulations were performed in PC/PG mixtures (1:1), but the production runs were completed in a mammalian-like mixture of PC/PS/Chol (6:2:2). To find the optimal conditions that would discriminate between the compressed and stretched states of the membrane, we started with the bilayer in an over-compressed state (∼48 Å² per lipid), then gradually expanded the simulation cell in the x and y dimensions, causing bilayer stretch. Analysis of the bilayer dynamics and the behavior of the adsorbed peptide during the 200 ns expansion ramp indicated that an area of 50.7 Å² per lipid represents the stable, tightly packed bilayer, while 65 Å² per lipid satisfies the state of the bilayer under lateral tension. In equilibrium simulations of these two states, we expected to observe significant changes in peptide position. The systems were equilibrated for 100 ns at each area per lipid, and production runs were continued for another 100 ns at constant area.

Figures 2A and 2B represent snapshots of representative peptide configurations in compressed and expanded bilayers, respectively, observed in 100-ns trajectories. The presented frames were chosen to be close to the average conformation along the trajectory. The conformations indicate that the peptide can change its orientation at the bilayer interface, simultaneously altering the depth of NBD insertion and the number of polar contacts (statistics are shown in Figure 3). Electrostatic maps of W6NBD in compressed and stretched bilayers (Figures 2C and D, respectively) depict positively charged peptides floating on top of the membrane slab with positive potential inside. This positive dipole potential is attributed to oriented water at the membrane interface (51, 52). It is, therefore, easy to imagine that the depth of insertion of the highly positively charged (+5) probe, besides lateral pressure, is limited by the common phospholipid interfacial dipole potential (50). This might also explain why our probes do not bind to pure zwitterionic PC membranes and require negatively charged lipids (below).

**Figure 2.**
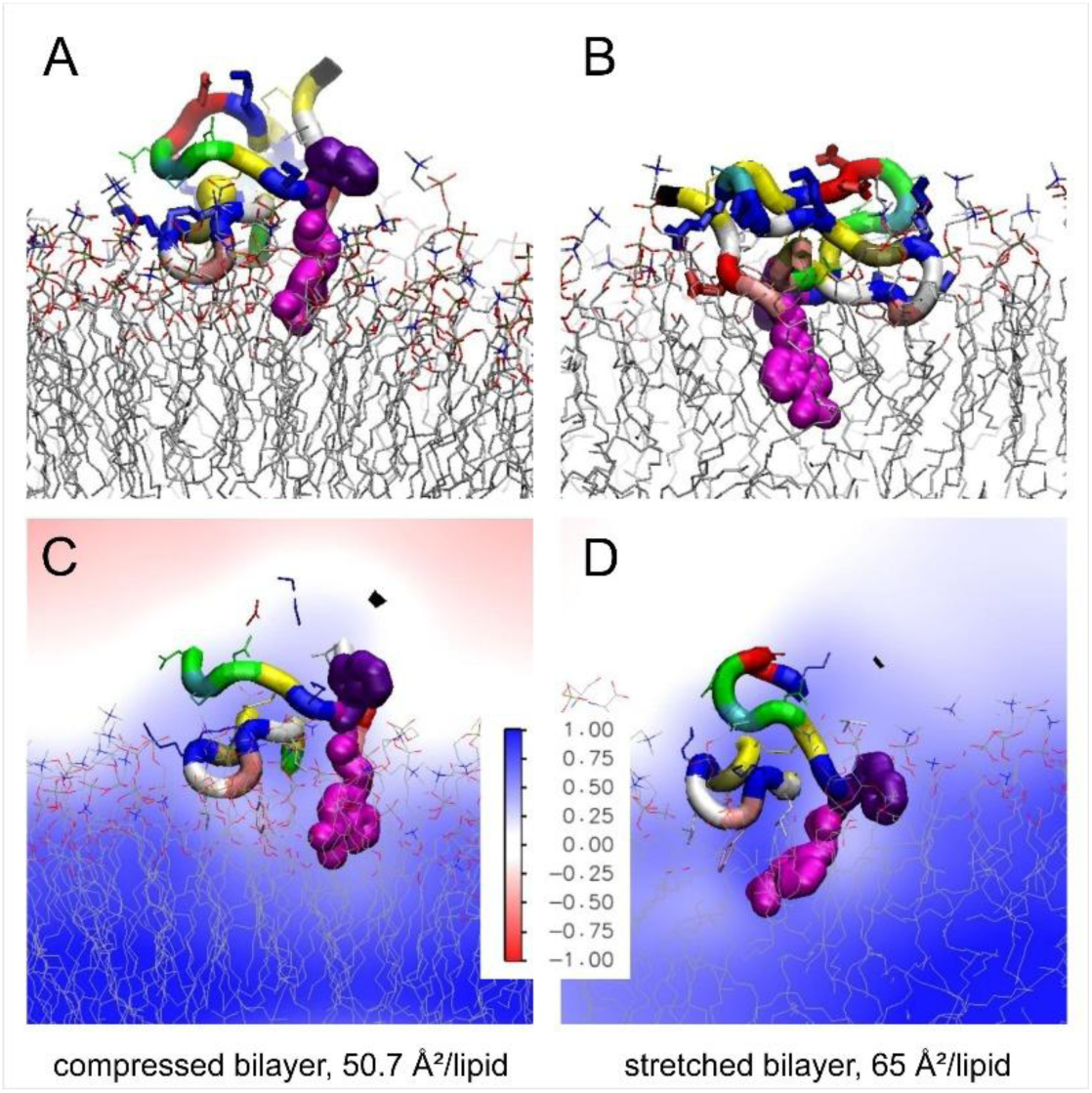
Representative conformations of the W6NBD peptide in simulated compressed (left) and stretched (right) lipid bilayers composed of PC/PS/Chol and the electrostatic maps of the system. Snapshots of the ‘shallow’ (A) and ‘deep’ (B) conformations illustrate the angular orientation of the peptide and the depth of the NBD group (purple) relative to the polar headgroup layer. The W7 sidechain near the interface is shown in dark purple. PME-calculated electrostatic maps of W6NBD peptides in compressed (C) and stretched bilayers (D). The color scale bar represents potentials from -1 to 1 V. The deep blue regions inside the membrane indicate the positive dipole potential at the membrane interface (50) interacting with the positively charged peptide.

**Figure 3.**
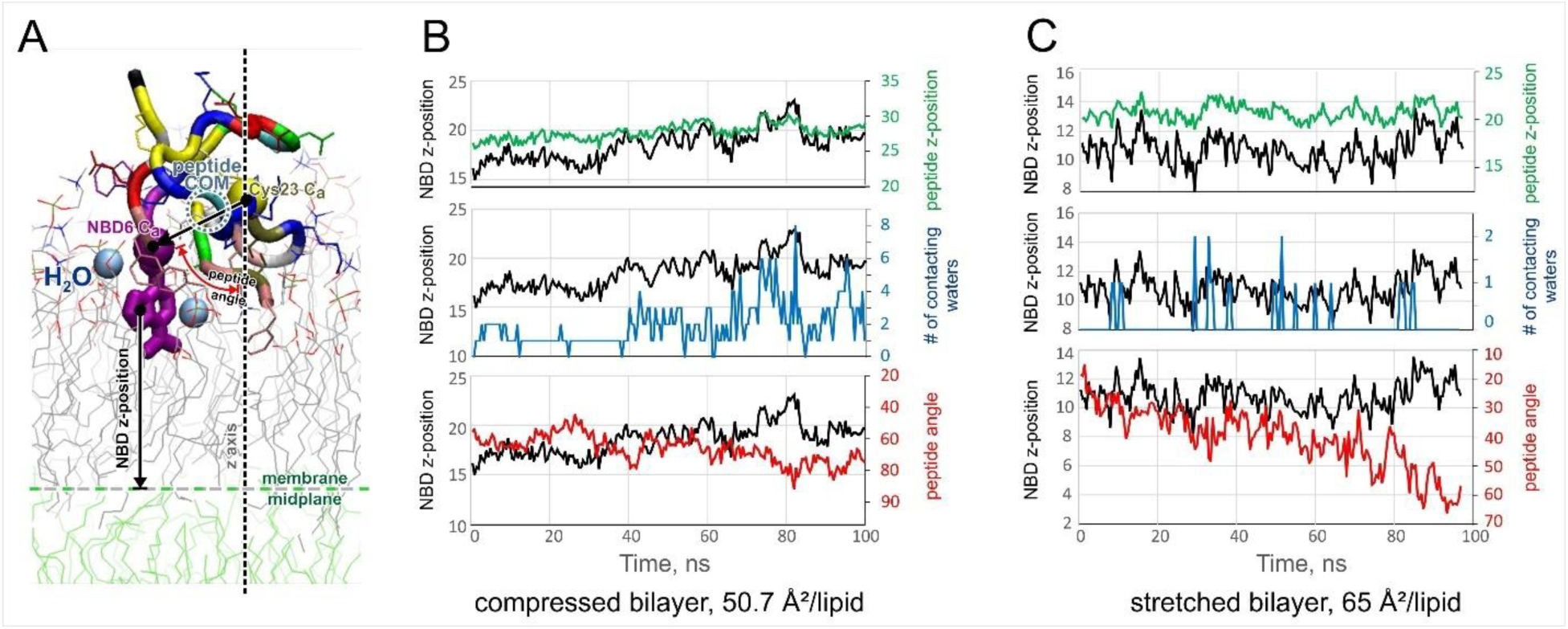
The statistics of simulated W6NBD peptide orientation, depth of insertion, and polar contacts of the NBD group in compressed and stretched bilayers. (A) The explanation of parameters extracted from 100-ns MD trajectories and shown as traces in panels (B) and (C): the distance of NBD from the midplane of the bilayer (black traces); the distance of the peptide’s center of mass from the midplane (green); the angle of peptide orientation (vector connecting Ca atoms of Cys23 and NBD amino acids relative to membrane normal); and the number of atomic contacts of NBD with water (blue). The transition from the shallow to the deep state involves about 4 Å deeper insertion and approximately a 60° change in the molecule’s orientation relative to the vertical z-axis.

The peptide’s conformation in the lipid bilayer, along with metrics characterizing its position and environment, is shown in Figure 3A. The parameters extracted from the trajectories (panels B and C) are: the angle between the vector connecting the alpha carbons of C23 and W6NBD amino acid and z-axis normal to the plane of the bilayer, the depth of the NBD group insertion into the monolayer, as well as the z-position of the center of mass of the peptide. We present trajectories as pairs to illustrate correlations between the depths of the NBD group (black traces) and the entire peptide insertion, the peptides’ angular orientation, and the number of polar water atoms in the immediate vicinity of NBD (the polarity of the environment). In the compressed bilayer, the peptide adopts a shallow conformation, positioning the fluorophore between polar groups. Its orientation fluctuates between 60 and 80 degrees from the normal (almost orthogonal to the z-axis), and the number of adjacent water molecules varies between 2 and 8 (Figure 3B). In the expanded (stretched) bilayer, the peptide stays closer to the upright position (30-40 degrees to the normal); the average position of the fluorophore changes as it moves 4-5 Å closer to the midplane of the bilayer, residing among aliphatic chains as illustrated by the low probability of NBD fluorophore contacts water (Figure 3C). By the end of the trajectory, the peptide rotated to ∼60 degrees, which negatively correlated with the depth of NBD insertion. The same insertion parameters for W7 are presented in Supplemental Figure S2, and the parameter statistics for both W6NBD and W7 sidechains in all four replicas are shown in Supplemental Table S1.

The trajectories indicate that the depth of the fluorophore insertion fluctuates within a range of 5-7 Å, with an approximate correlation time of 5-10 ns. The simulations demonstrate the scale of the transition between the shallow and deeply inserted states in a bilayer mimicking a mammalian membrane with PS as the main anionic lipid species. The experiments presented below also examine phosphatidic acid (PA) and phosphatidylglycerol (PG) as two other anionic species.

## Langmuir monolayer experiments illustrate insertion and pressure-dependent redistribution of W6NBD at the monolayer-bilayer equivalence pressures

Monolayers represent half-membranes (53). This method allows us to vary the lateral pressure of lipids over a wide range while controlling the area, measuring the lipid film’s compressibility, and determining the areal contribution of lipid-intercalating molecules. In this standard setup, we also measure fluorescence from a monolayer during compression and typically observe the following sequence of events. At the start of an experiment (excess surface area with wide-open barriers, zero lateral pressure), the peptides and lipids exist as a mixed two-dimensional gas at the air-water interface. As barriers close, the lipid-peptide mixture forms a continuous phase, and both pressure and fluorescence initially increase with monolayer compression, reflecting higher lipid and probe densities. Then, the monolayer’s stiffness and fluorescence both exhibit peaks, followed by decreases as compression continues. The correlated drop in these parameters reflects the onset of peptide expulsion from the monolayer into the subphase or into the shallow adsorbed state. With GsMTx4-based peptides, this transition uniquely occurs in the region around the monolayer-bilayer equivalence pressure, estimated between 32 and 41 mN/m for mammalian-type lipids (54–56). At this pressure, the lipid packing density in the monolayer matches that of an unstressed bilayer. In this state, the peptide probe is predicted to be primarily in the ‘shallow’ low-fluorescence adsorption state. A monolayer in its expanded state corresponds to the highly stretched membrane where the probe tends to insert deeply. Therefore, probe expulsion under lateral monolayer pressure demonstrates the probe’s insertion into the stretched bilayer, only in reverse. Since monolayer compression isotherms are not fully reversible, this method works in only one direction, allowing the observation of expulsion as if the membrane’s tension is being released. The relationship between the tension-driven incorporation of fluorescent probes into the stretched membrane and pressure-dependent expulsion from the monolayer is illustrated in Supplemental Fig. S3.

Our first experiment examined how much the probe incorporates into a relaxed monolayer and how much remains in the subphase. This is illustrated in Supplemental Figure S4 and its legend. The laser beam was aimed at a spot near the edge of the monolayer trough, illuminating the area that the barrier would pass during compression. The probe was dissolved in the subphase buffer at 0.3 µM, and the aqueous probe’s fluorescence was measured before applying the lipid. After spreading the lipid, the probe entered the monolayer over a 30-minute time course. Then, the barriers began to move, causing compression, and when one barrier completely crossed the beam, sweeping the lipids to the middle of the trough, fluorescence dropped to about 5% of the initial value. This indicates that 95% of the probe was incorporated into the lipid film. For a trough depth of 0.1 cm and a typical diffusion coefficient of 3×10^-6^ cm^2^/s for small proteins, the incorporation time is estimated to be approximately 30 minutes. Here, we clearly encountered a diffusion limitation for peptide binding and insertion into the lipid interface.

We investigated several monolayer compositions containing phosphatidylcholine (PC) as the base lipid and three different types of anionic lipids: phosphatidylserine (PS), phosphatidic acid (PA), and phosphatidylglycerol (PG), which serve as electrostatic attractors for the positively charged NBD peptide. PG, a less common lipid in mammalian plasma membranes, also demonstrated a strong affinity for attracting W6NBD into the deep adsorption state in liposomes without tension (see below). For this reason, we paid less attention to PG as an electrostatic attractor; the monolayer data for two PC/PG mixtures are presented in Supplemental Figure S5. We also studied the effects of lipid-ordering additives such as sphingomyelin (SM) and cholesterol (Chol). We began with the mammalian-type mixtures of PC, PS, SM, and Chol.

Figure 4 illustrates the behavior of W6NBD with monolayers of three compositions: PC/PS/Chol (6:2:2, column A), PC/PS/SM (6:2:2, column B), and PC/PS/SM/Chol (6:2:1:1, column C). The upper row shows isotherms recorded in the presence of 0.3 µM W6NBD, compared with control compression isotherms (gray). The peptide ‘swells’ monolayers, shifting the isotherm up and to the right, meaning that it incorporates and takes up a substantial area in the lipid film. At higher surface pressures, the two isotherms converge, indicating that lateral pressure expels the peptide back to the subphase or to the shallower adsorption state.

**Figure 4.**
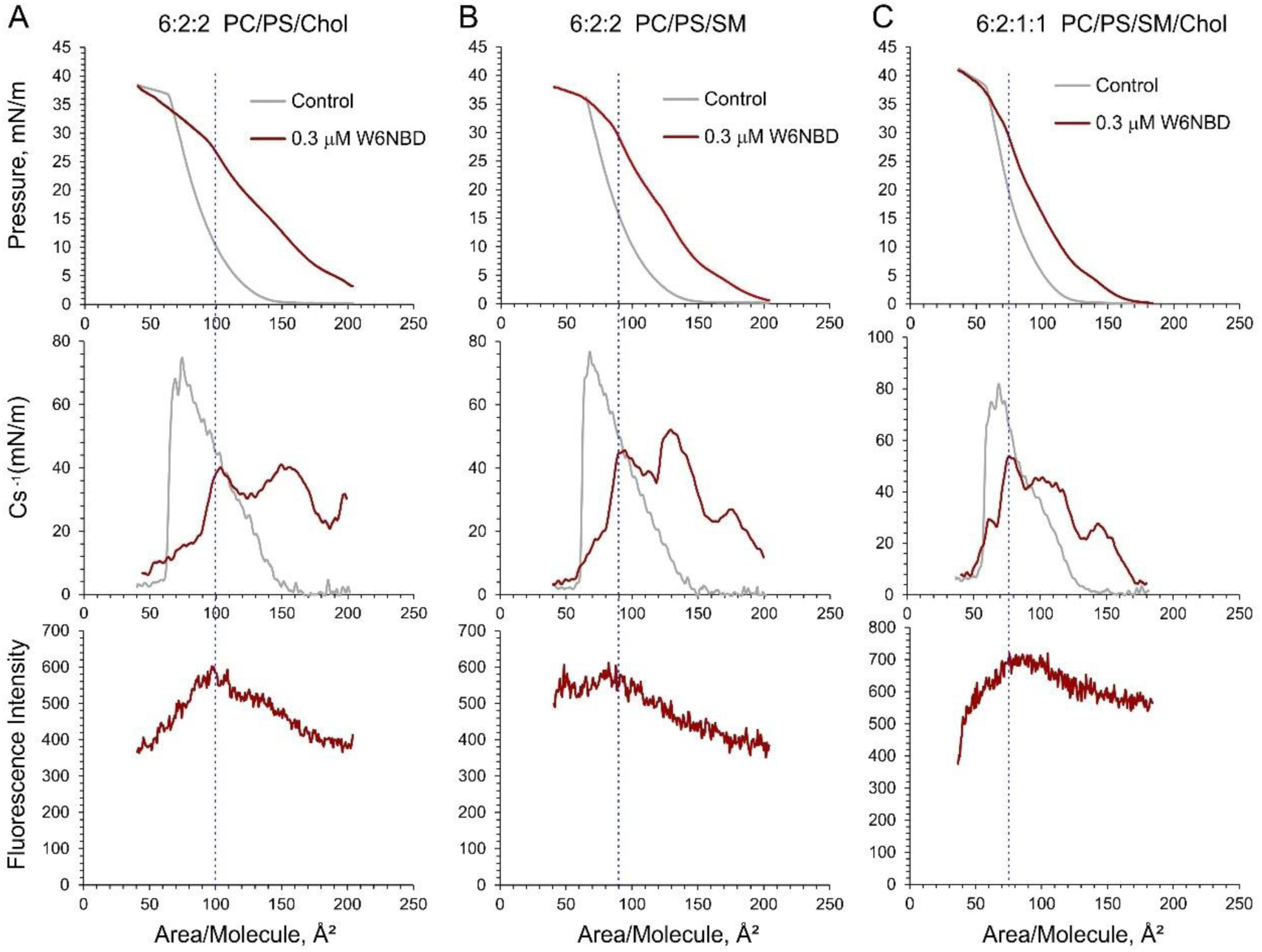
Compression isotherms for three PC/PS/SM/Chol mixtures recorded in the presence of 0.3 µM W6NBD in the subphase. W6NBD was allowed to equilibrate for 30 min. Gray traces represent controls without the peptide. The fluorescent peptide was allowed to enter the newly formed monolayer for 30 min. The upper row shows the initial compression isotherms, which bend in the presence of W6NBD at pressures between 27 and 30 mN/m. The plots of inverse compressibility modulus Cs^-1^ = A(dπ/dA) amplify the irregularity of the isotherm slope (middle row). The peak of Cs^-1^, marked by vertical dotted lines, and the following drop reflect the expulsion of the peptide from the monolayer, which coincides with the drop in fluorescence (bottom row) as the compression proceeds.

The isotherms exhibit breaks in slope and inflections, visualized as peaks and troughs on the inverse compressibility modulus plots, which represent stiffness (Cs^-1^). The left-most peak of Cs^-1^, indicated by the vertical dotted line, marks the beginning of peptide expulsion, which precisely aligns with the drop in fluorescence (bottom row). The fluorescence traces show that at the beginning of compression (at the right end), the probe is rarified in the expanded monolayer; as compression proceeds, it condenses with lipids, reaches a peak, and then is gradually expelled. The simultaneous drop of stiffness and fluorescence indicates the re-partitioning of the probe to a more polar environment. In the PC/PS/Chol mixture, expulsion begins at ∼28 mN/m and continues beyond the intersection point with the control isotherm at 35 mN/m. The expulsion phase covers a ∼10 mN/m range around the monolayer-bilayer equivalence pressure (∼35 mN/m). The PC/PS/SM mixture exhibits a steeper slope of the isotherm, but also a smaller decrease in fluorescence from peak to collapse. The quaternary mixture (PC/PS/SM/Chol) exhibits an even steeper slope, indicating that the presence of Chol and SM creates a less accommodating environment for the peptide. However, based on fluorescence, SM alone (column B) provides stronger probe retention up to monolayer collapse. In contrast, the addition of Chol on top of SM (column C) enables expulsion, albeit at slightly higher lateral pressure and lower area per lipid.

We observed a similar compression-dependent behavior of W6NBD when PA was chosen as the electrostatic ‘attractor’ for the peptide (Figure 5). The left column illustrates the swelling of the expanded PC/PA/Chol monolayer with the incorporation of the W6NBD peptide, followed by expulsion. In the PC/PA/Chol mixture, the expulsion begins at approximately 28 mN/m and continues to the point of collapse, converging with the control isotherm at 40 mN/m, thus covering the 12 mN/m range around the monolayer-bilayer equivalence pressure (∼35 mN/m). The Cs^-1^ and fluorescence plots show perfect coincidence of compressibility and fluorescence changes, marking the onset of expulsion.

**Figure 5.**
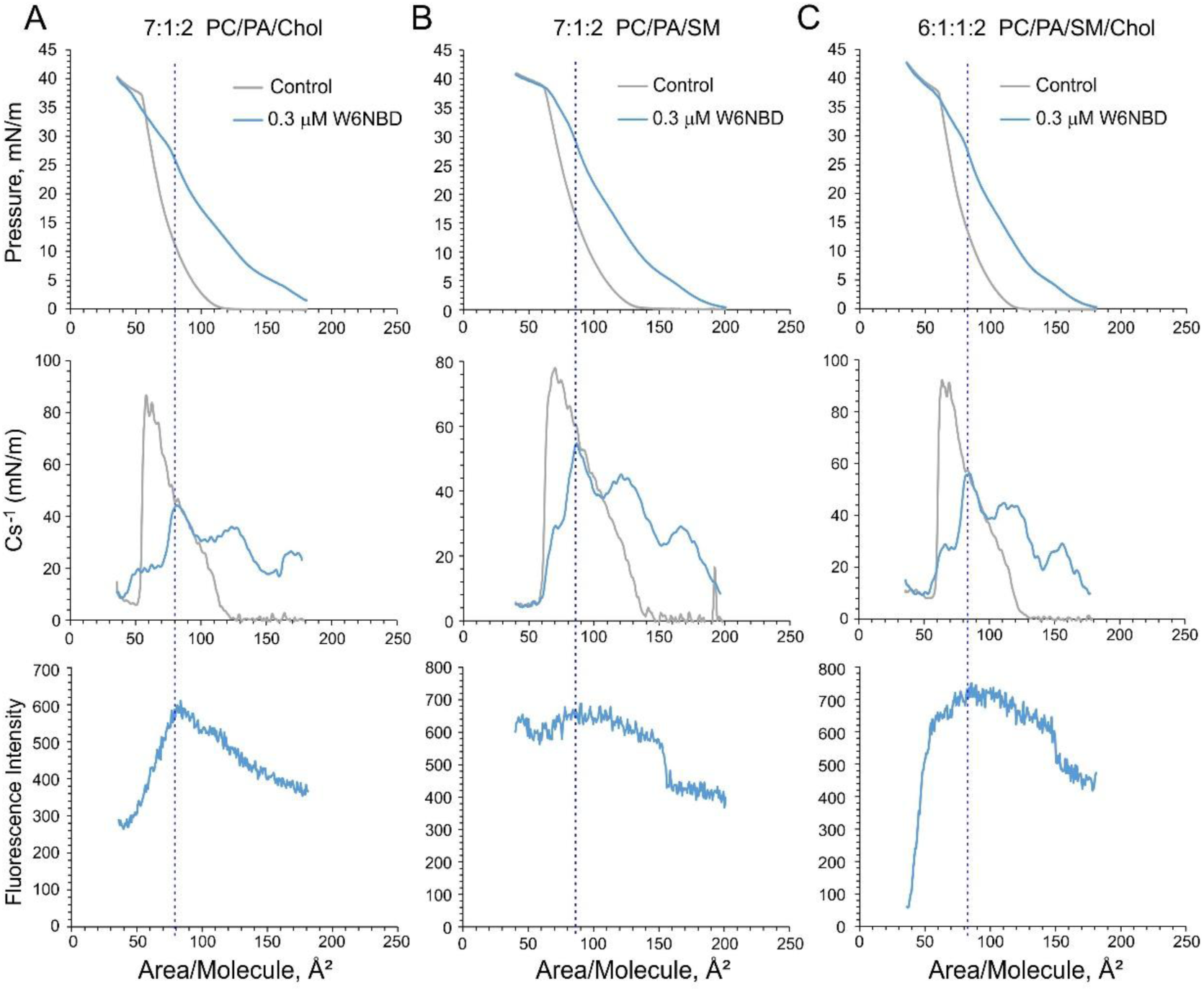
Compression isotherms for ternary PC/PA/Chol (A), PC/PA/SM (B), and the quaternary PC/PA/SM/Chol (C) mixtures with incorporated W6NBD. Gray traces represent controls without the peptide; blue traces represent experiments with 3×10^-7^ M W6NBD in the subphase. After spreading the lipid, the fluorescent peptide was allowed to incorporate for 30 min. The upper row represents the original compression isotherms, which visibly bend in the presence of W6NBD at pressures between 27 and 30 mN/m. Peaks in the plots of inverted compression modulus Cs^-1^ (second row) show this irregularity of the slope. The drop of Cs^-1^ at this point reflects an increase in compressibility related to the beginning of the expulsion of the peptide from the monolayer. This expulsion is confirmed by the simultaneous drop in fluorescence (bottom row) as the compression proceeds.

The isotherms in the middle column (B) show that adding SM instead of Chol alters the partition-pressure curve and the shape of Cs^-1^ peaks. It slightly raises the expulsion threshold (to ∼32 mN/m) and, as previously, shows a greater retention of W6NBD in the monolayer, as indicated by the fluorescence curve that does not sharply decline to the left of the peak. When all four components, including SM and Chol, are present (column C), they further modify the Cs^-1^ peaks but allow the probe’s expulsion at 28 mN/m. The shape of the fluorescence curve indicates that the most significant redistribution of the probe to the shallow state/subphase occurs at higher pressures and smaller lipid area.

With both electrostatic attractors, PS or PA, the two ‘ordering’ components, SM and Chol, produce distinct effects: SM increases the retention of the probe, while Chol promotes its expulsion. The two ‘ordering’ components slightly alter the pressure expulsion threshold (28-32 mN/m). Columns C in both figures, representing quaternary mixtures with SM and Chol, suggest that these bilayers will resist deep probe incorporation into the resting membrane but will allow incorporation into expanded bilayers under tension.

The isotherms and the Cs^-1^ curve recorded in the presence of SM clearly indicate probe expulsion, but the fluorescence trace lags behind and does not drop significantly. This result suggests that SM may result in a different ‘expelled’ state (a shallow bound position) of the probe, which might still have its NBD group submerged in a less polar environment.

Regarding the difference between PS and PA as electrostatic attractors, the isotherm break for the PC/PS/Chol mixture (Figure 4, column A) occurs at a higher area per molecule with a shallower slope between the break and monolayer collapse. The PC/PA/SM displays a steeper slope (Figure 5, column A), and the quaternary mixture is even steeper. While PS carries a phosphate in a diester configuration and a carboxyl, PA has a compact phosphomonoester headgroup and, when interacting with lysines on our peptides, may additionally deprotonate and carry a -2 charge. Thus, PA may attract the W6NBD peptide more strongly than PS.

A clear difference between membrane-ordering components, SM and Chol, was evident from the analysis of W6NBD expulsion rates with pressure, which directly reflects effective molecular areas of intercalation (Supplemental Figure S6). It is easy to see that the excess area of the monolayer caused by the ‘guest’ peptide reflects the mole fraction of the ‘guest’. The slope of the decrease in monolayer area with increasing lateral pressure provides an estimate of the area occupied by individual ‘guest’ molecules within the monolayer plane. The slopes and deduced areas clearly depend on the type of lipid-ordering components present, with the lowest (∼110 Å²) observed in the PC:PA:SM mixture and the highest (∼250 Å²) recorded in the PC:PA:Chol mixture.

## The incorporation of peptides into unperturbed and osmotically shocked liposomes exhibits a strong electrostatic component

Standard fluorometry was used to test the incorporation of the peptide into liposomes of various compositions (Fig. 6). To make unilamellar liposomes osmotically sensitive, they were extruded through a 0.2 μm nucleopore filter in a 10 mM phosphate, 100 mM NaCl buffer supplemented with 0.5 M sorbitol, resulting in an osmolality of 0.7 Osm, both inside and outside. W6NBD was used uniformly at a concentration of 0.3 μM; its fluorescence level in pure buffer was set as unity for normalizing all fluorescence changes. Mixing liposomes with W6NBD increased fluorescence, indicating that the adsorbed peptide inserts into the lipid bilayer, and some NBD fluorophores now reside in a nonpolar environment. The exception was pure POPC liposomes, which showed no detectable change in fluorescence, suggesting either no affinity for W6NBD or very shallow adsorption (data not shown). Adding negatively charged lipids such as PA, PG, or PS significantly alters the binding behavior, as seen by fluorescence levels that increase with the mole fraction of the anionic species (Figure 6E). At moderate mole fractions, these anionic lipids, acting as electrostatic attractors, also cause a greater fluorescence increase due to osmotically generated tension. At higher mole fractions, these anionic lipids saturate W6NBD binding to unperturbed membranes, eliminating responses to tension, as exemplified by the PC/PG mixture (Figure 6A). Our observations indicate that PA is the most effective electrostatic attractor for W6NBD, PS has intermediate potency, and PG strongly promotes W6NBD binding to the ‘deep’ membrane position, thereby abolishing the tension-dependent increase.

**Figure 6.**
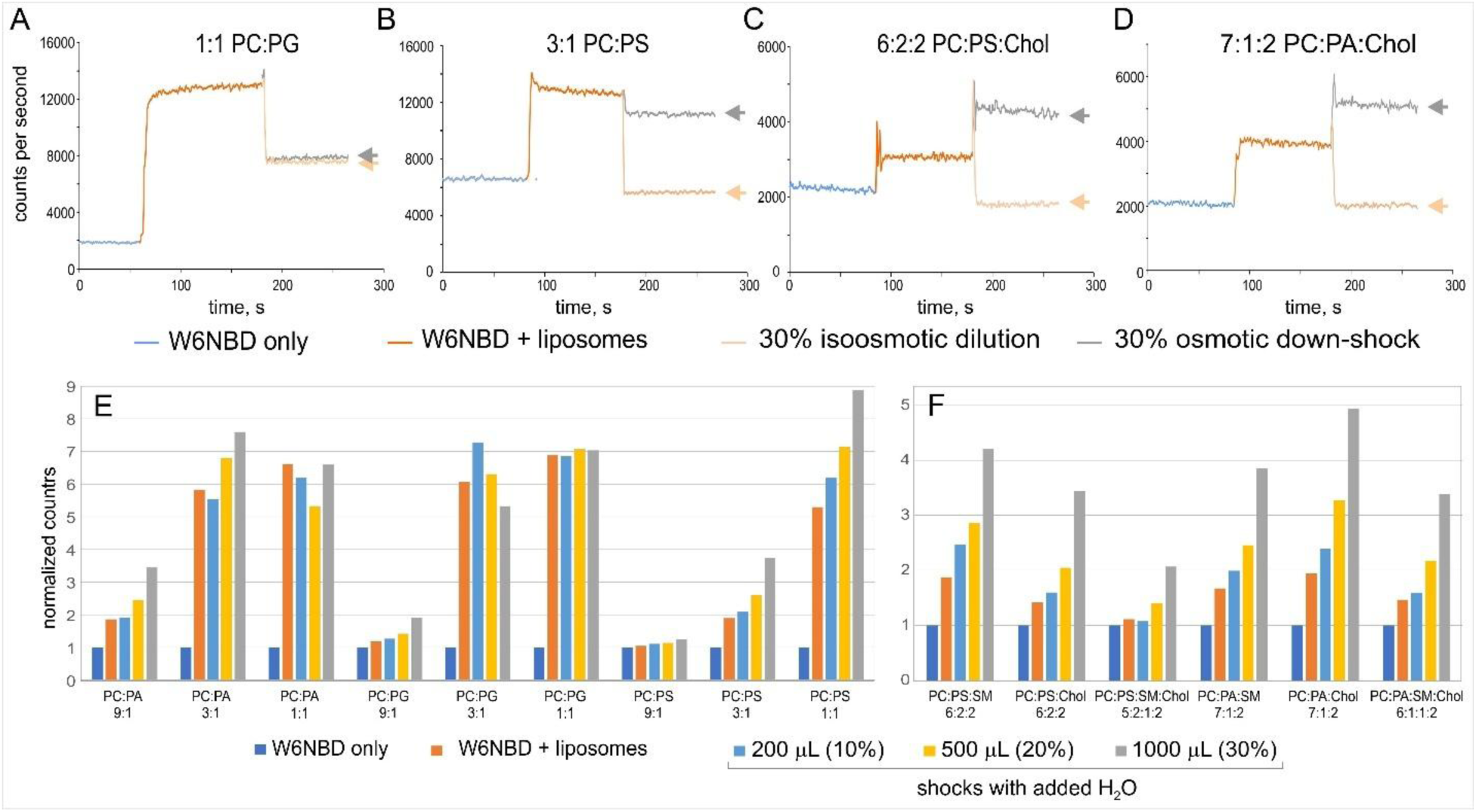
Effects of lipid composition on the affinity of W6NBD to the liposomes and osmotic shock-induced fluorescence change. (A) Fluorescence levels (counts per second) upon sequential addition of W6NBD to the cell, liposomes, and then either dilution with 1 mL of isosmotic buffer (beige arrows) or 1 mL of distilled water (osmotic downshock, grey arrows). For PC/PG liposomes, dilution and osmotic shock produced identical effects. PC/PS (B), PC/PS/Chol (C), and PC/PA/Chol (D) liposomes exhibited a 3- to 5-fold increase in fluorescence upon osmotic downshock. (E) Effects of the mole fraction of three anionic lipids (PA, PG, and PS) on W6NBD incorporation into liposomes prior to and upon osmotic shock. (F) Effects of lipid-ordering components, Chol and sphingomyelin SM, on W6NBD incorporation into PC/PA or PC/PS mixtures. These two components reduce background insertion and effectively increase the fold change in fluorescence upon osmotic shock. Steady-state fluorescence was recorded over 60-80 s, then 0.2 μm liposomes of the desired composition were added, and the recording continued for an additional 100 s. The initial increase in fluorescence was associated with the insertion process into osmotically unperturbed liposomes. The levels after the osmotic shock or isosmotic dilution are shown by gray and beige arrows in (A), and the ratio of these levels reflects the effect of osmotic stress. The pre-shock fluorescence level (counts per second) was multiplied by this ratio to quantify the effect of osmotic stress. In the bar graphs, all data are normalized to the fluorescence level of free W6NBD in the buffer. The bars represent averages from 3 independent experiments, with the standard deviation of normalized counts within 6%.

## Membrane-ordering components diminish resting incorporation of W6NBD, but increase the dynamic range of tension responses

After selecting an appropriate anionic component, we examined mammalian-like compositions that include membrane-ordering components such as Chol and/or SM. As shown in Figures 6C and D, adding Chol to PA- and PS-containing liposomes reduced the W6NBD fluorescence upon liposome injection by 10-12 fold compared to PC/PA or PC/PS mixtures. This indicates tighter lipid packing that resists deep penetration into the unstressed bilayer. At this low baseline of resting fluorescence, osmotic shock causes a more gradual increase in fluorescence than negative lipids alone (Fig. 6F). This behavior of the fluorescent probe clearly indicates that tension alters membrane packing order, allowing the peptide to penetrate more deeply. In line with the monolayer experiments (Fig. 3 and 4), SM has a milder ordering effect, allowing slightly more probe to incorporate into both PA- and PS-containing membranes at rest, whereas Chol results in a stronger exclusion of W6NBD from unstressed membranes. The behavior of quaternary mixture 6:1:1:2 PC/PA/SM/Chol was intermediate between ternary mixtures containing either SM or Chol. The quaternary mixture of 5:2:1:2 PC/PS/SM/Chol showed dominance of Chol, with almost no incorporation at rest and only a moderate 2-fold increase in fluorescence upon a strong osmotic shock. These manual-injection steady-state fluorescence measurements identified preferred compositions suitable for testing in time-resolved kinetic experiments.

### Stopped-flow osmotic shock experiments reveal the kinetics of stretch-dependent probe insertion

To analyze the kinetics of SUV swelling and W6NBD incorporation, we used a two-syringe stopped-flow system (SFM2000, BioLogic). The state of the W6NBD probe was monitored under a 460 nm laser excitation, and the intensity of a broad emission band (> 525 nm) was collected. Syringe 1 contained 0.3 µM W6NBD and 50 µg/mL of 0.2 µm extruded liposomes suspended in a sorbitol-containing buffer (0.685 Osm). Syringe 2 held distilled water. The contents of both syringes were mixed to a total volume of 302 µL; volume ratios ranged from 1:3 to 6:1, resulting in a final osmolarity ranging from 627 to 183 mOsm (Figure 7). Reproducible fluorescence changes were observed at osmotic downshifts greater than 58 mOsm (685 → 627).

**Figure 7.**
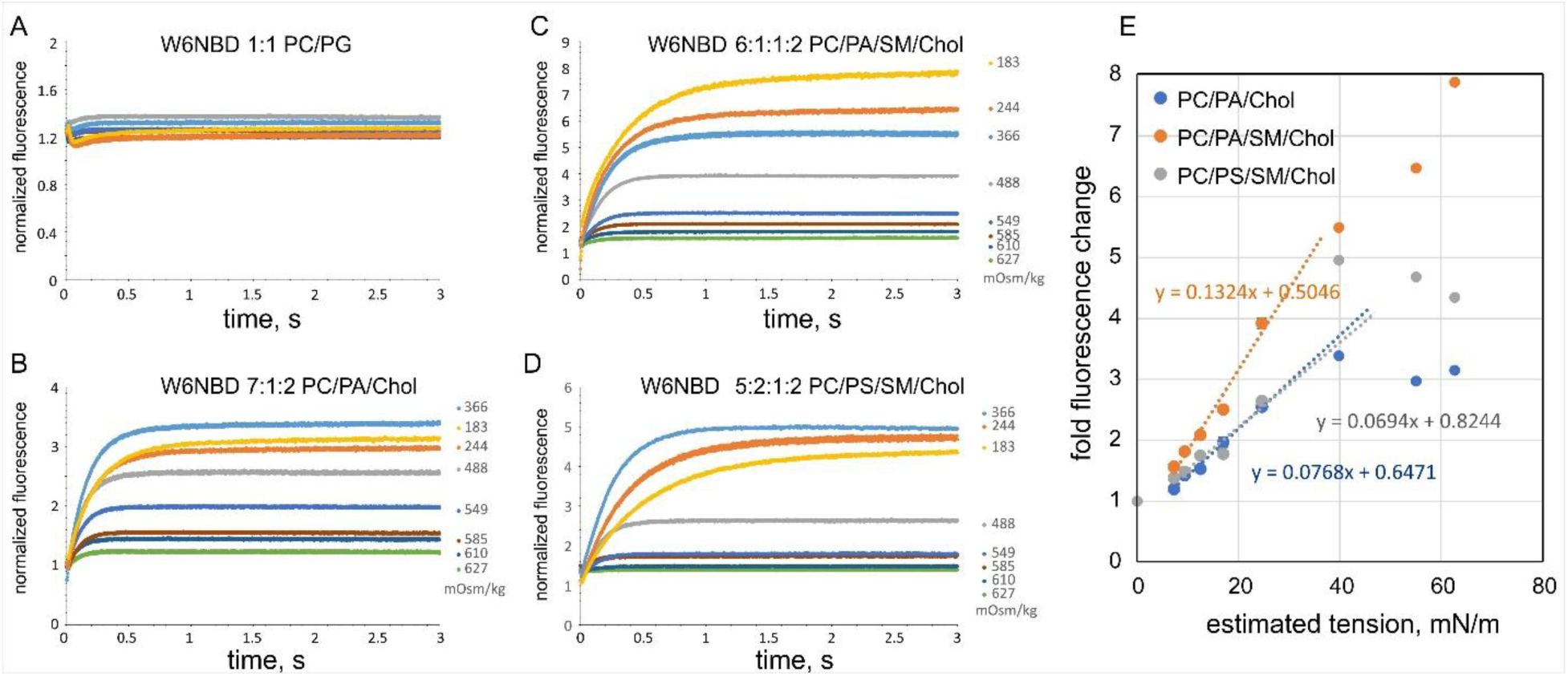
Stopped-flow osmotic shock experiments track the kinetics of W6NBD incorporation and its transition to the deep, high-fluorescence state in liposomes of varying composition. The liposome compositions are shown above the traces. (A) PC/PG liposomes showed virtually no response to osmotic stress. (B) PC/PA/Chol liposomes exhibited about a 3.3-fold increase, saturating at 319 mOsm downshift. (C) The PC/PA/SM/Chol mixture shows the steepest response in the shock range and a greater fold increase in fluorescence at extreme shock levels. (D) The PC/PS/SM/Chol mixture shows an earlier ‘beak’ in the rising fluorescence versus shock magnitude and a lower slope. In all experiments, the background, recorded from pure buffer with W6NBD (∼5% of the signal), was subtracted, and the fluorescence traces were normalized at appropriate dilution ratios to the unshocked controls, where liposomes were mixed with the 685 mOsm sorbitol buffer. The steady fluorescence level at the end of the stopped-flow traces was plotted as a function of the estimated tension in panel E. The hydrostatic pressure p inside 0.2 μm liposomes, generated by a drop in external osmolarity, was calculated using the Van’t Hoff equation (p = RTΔC), and tension was estimated using the Young-Laplace relation for a spherical liposome (γ = pr/2).

In stopped-flow experiments, we observed a similar dependence of fluorescent signals on liposome composition as in the non-timed osmotic shock experiments above. Upon mixing with W6NBD, the PC/PG liposomes exhibited high fluorescence right from the start. The osmotic downshock did not significantly change their fluorescence levels (Figure 7A). We then tested compositions identified in previous experiments as more responsive to osmotic stress. Figure 7B shows the kinetics of W6NBD osmotic shock responses for PC/PA/Chol liposomes, displaying up to a 3.3-fold increase in fluorescence with a 319 (685 → 366) mOsm osmolarity decrease. We estimate that this osmotic gradient could potentially produce a membrane tension of 38 mN/m in a liposome with a diameter of 0.2 μm. We emphasize that this is just an estimated tension, and it is unlikely that the liposome can withstand this stress for an extended period. As shown in the graph, increasing the shock magnitude beyond a certain point did not further increase the fluorescence level, indicating that the liposomes ruptured and became leaky at those magnitudes. Liposomes made of a PC/PA/SM/Chol mixture (panel C) showed nearly linear growth in response (fluorescence end level) as a function of the shock magnitude, indicating that this composition forms more stable membranes. The PC/PS/SM/Chol membranes (panel D) were somewhat less stable than the quaternary mixture with PA, but they still produced a fivefold increase in fluorescence, peaking at 319 (685 → 366) mOsm osmolarity drop.

Based on the slope of fitting curves in Figure 7E, liposomes made with a ternary PC/PA/Chol mixture respond to a decrease in external osmolarity with a nearly linear fluorescence increase of about 7% per mN/m. The quaternary PC/PA/SM/Chol mixture exhibits a steeper fluorescence response, with approximately 13% increase for each mN/m in the tension range between 6 and 22 mN/m. The quaternary PC/PS/SM/Chol mixture produces responses with a lower slope, corresponding to roughly 6.7% fluorescence increase per mN/m within the same tension range. Except for the quaternary PC/PA/SM/Chol mixture, which shows a steady increase beyond an estimated 30 mN/m, the tension responses of the other mixtures tend to bend downward, indicating liposome osmotic rupture.

Exponential fits of the traces revealed different time constants depending on the magnitude of the downshock. Small downshocks (685→627 mOsm) showed an equilibration time of 80-100 ms, intermediate shocks (685→549 mOsm) had tau values of 110-130 ms, and extreme shocks resulted in longer times, ranging from 240 to 480 ms. This seems to be related to the fold of dilution of free W6NBD upon mixing the liposome suspension with distilled water at different ratios. The stretched liposome membranes have a higher affinity for the probe, and the slower onset of fluorescence reflects the longer diffusion time to the liposome surface.

### Pipette aspiration of GUVs in the presence of W6NBD and W7NBD peptides

To determine the rapid dynamics and reversibility of the tension-dependent responses of NBD peptides, we measured fluorescence changes in giant unilamellar vesicles (GUVs) generated using electroformation (57). To investigate how the fluorescence response depends on the fluorophore’s location on the peptide, we compared peptides with NBD substitutions at either W6 or W7. MD simulations suggested that W7 has a shallower resting position than W6NBD (Supplemental Figure S2), but the higher sensitivity of imaging the bilayer only with an EM CCD camera allowed us to assess the W7 response. Tension was applied using pipette aspiration (58–60) in the presence of W6NBD or W7NBD. Both peptides exhibited a strong increase in fluorescence with increasing pressure, which depended on lipid composition (Figure 8A and 8B). A series of pressure pulses with gradually increasing amplitudes in -3 mmHg steps was used to create the relations shown in Fig. 8F-G (See supplementary video 1 for W6NBD response). Each pulse lasted 5 seconds, giving the vesicle time to reach a steady state, followed by 20-second periods during which the pressure was returned to 0 mmHg (see Supplemental Video 1) to assess reversibility. The rupture resistance of GUVs to tension was significantly lower than that of SUVs, possibly due to the larger radius of curvature and uneven curvature stress at the pipette opening. Similar to earlier experiments reported by Rawicz (58), we reliably achieved tensions up to approximately 6 mN/m. Within this range, we observed a similar pattern of lipid-dependent resting fluorescence and tension sensitivities as seen with the SUVs for both peptides. As a negative control, we used FM1-43, which in our experiments behaved as a completely tension-insensitive dye, in stark contrast to our probes, W6NBD and W7NBD.

**Figure 8.**
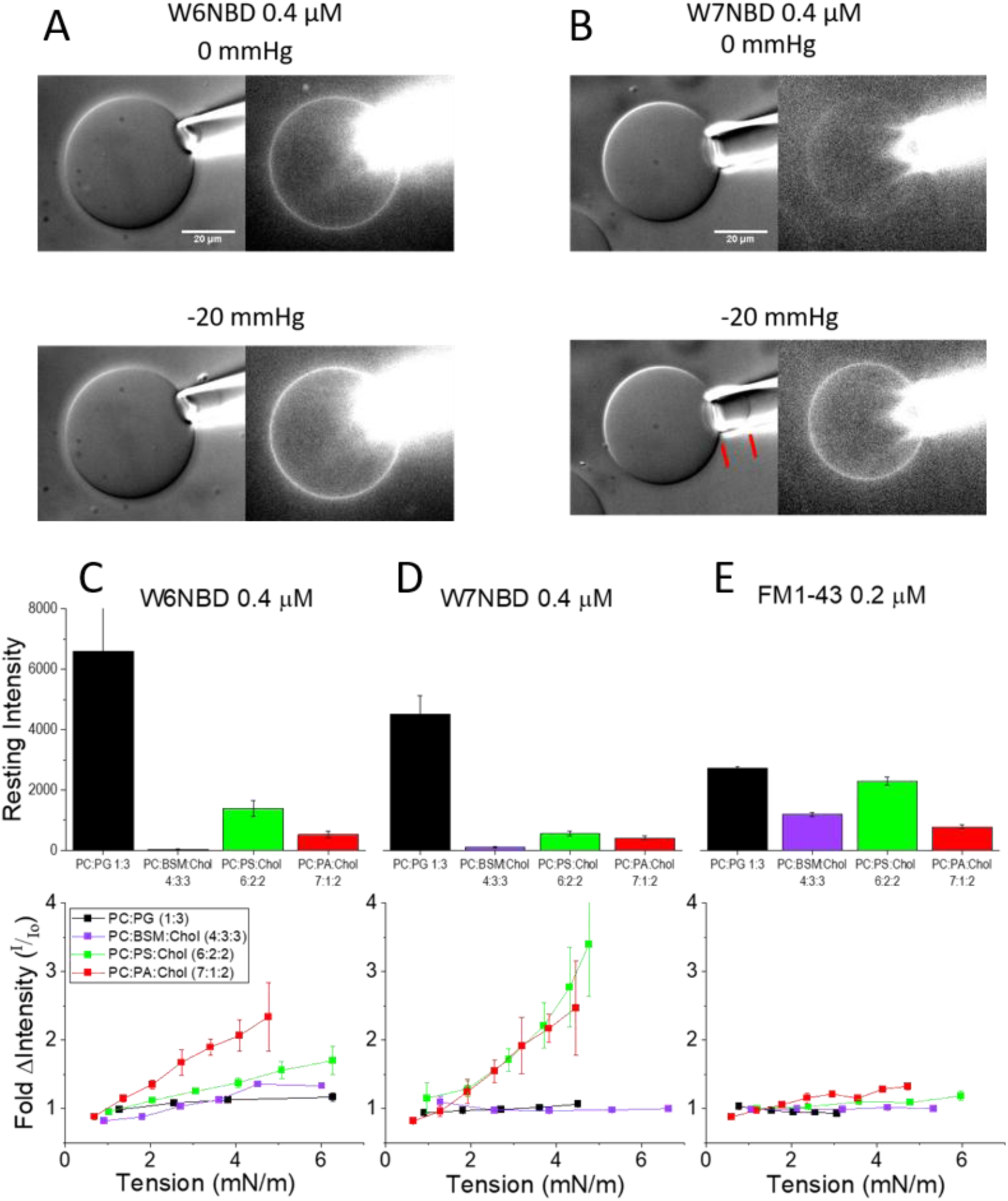
Steady state fluorescence measurements in GUVs. Vesicle aspiration experiment showing DIC and 520 nm fluorescence images of 6PC:2PS:2Chol vesicles treated with either W6NBD (A) or W7NBD (B) at rest and under -20 mmHg pressure. While fluorescence from the peptide bound to the glass pipette obscured the vesicle protrusion length, this distance could be easily measured in the DIC images, as indicated by the red lines in B. The resting fluorescence of W6NBD, W7NBD, and FM1-43 (control) in different vesicle compositions is shown in top panels of C-E. The vesicles were aspirated to different pressures, and the fluorescence change was allowed to reach a steady state over 4-5 seconds. After each pressure change, vesicles were allowed to relax at 0 mmHg pressure before a subsequent larger pressure step was applied. The tension change was calculated based on the aspirated protrusion length and pressure, as described by Rawicz (58). The change in fluorescence intensity was plotted with respect to the calculated tension in C-E lower panels.

As in the SUV experiments, GUVs composed solely of PC and the ordering agents Chol and SM showed negligible fluorescence in the relaxed state. In contrast, negatively charged PC:PG vesicles displayed very high resting intensity, representing the extreme conditions (Fig. 8C and D, upper panels). Correspondingly, both peptides showed no response to tension in these lipids within the 6 mN/m range (Fig. 8C and D, lower panels). However, when PS or PA was used as an electrostatic attractor alongside PC and Chol, we observed significantly lower resting background fluorescence with W6NBD than with PG/PC mixtures. W6NBD demonstrated strong tension sensitivity, with PA-containing vesicles (∼33% change/mN/m) nearly doubling the response seen in PS (∼15% change/mN/m), consistent with the monolayer and SUV data. Although the % change in fluorescence relative to tension was much higher in GUVs than in SUVs, the lipid sensitivities were proportionally similar. The lower % fluorescence change in SUVs may be due to deeper resting penetration depths, since the curvature at ∼100 nm radius in SUVs could affect peptide penetration at rest. Higher tension sensitivity seems to depend on a low initial resting fluorescence, as PS-containing vesicles nearly doubled the resting intensity compared to PA vesicles. Supporting this, W7NBD showed a similar tension response to both PA and PS and had comparable low resting intensities in both compositions. W7NBD displayed a tension response similar to that of W6NBD, with approximately a 33% change/mN/m, and was slightly greater in PS-containing vesicles. It also showed no tension sensitivity in either PC:PG (where the resting fluorescence saturated deeply) or PC + ordering agents (lacking negative attractors and with high resting stiffness). As a control, the membrane marker FM1-43, which inserts deeply into the membrane at rest, exhibited little dependence of resting intensity on lipid composition and no tension sensitivity (Fig. 8E, upper and lower panels respectively).

To assess the dynamics of the tension-dependent transition, we measured the rate of fluorescence change at 40 frames per second for both W6NBD and W7NBD peptides in response to 1.5 sec pressure steps of varying magnitude (Fig. 9A, and supplementary video 2 for W7NBD response). Surprisingly, the combination of charged and ordering lipids (PG:SM:Chol) without PC produced one of the most robust and reproducible responses to a programmed series of tension pulses. The fluorescence responses are plotted relative to the resting intensity for W6NBD (Fig. 9B) and W7NBD (Fig. 9C). The relative response of W7NBD was proportionately larger (30% change at 40 mmHg) compared to W6NBD (10% change at 40 mmHg). This difference may reflect the more than 10-fold lower relaxed intensity of the W7NBD, indicating that more NBD moieties occupy the shallow position primed for rapid transition. Notably, the intensity changes for both analogs were biphasic, featuring a rapid increase closely following the pressure change (τ < 100 ms), followed by a slower increase with τ of 0.5-1 sec. The relaxation response to tension release exhibited a similar two-phase decrease in fluorescence. The fast phase may represent a rapid transition of bound peptides from shallow to deep occupancy, while the slower phase reflects the association of new peptides from the bulk or the reorientation of peptides on the surface. Both peptides had similar kinetics.

**Figure 9.**
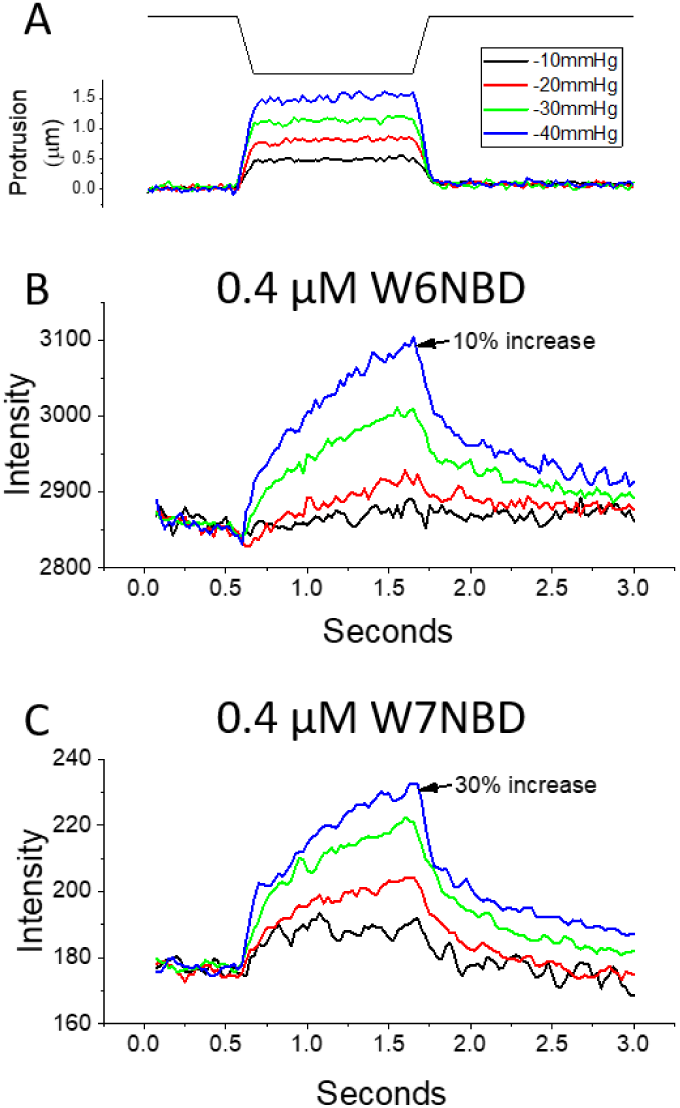
High (25 ms/frame) temporal resolution of dynamic fluorescence response. Mechanical stimulation of aspirated ternary 4PG:3SM:3Chol vesicles was produced with trapezoidal 1.5 s pressure pulses (80 ms rise time) of varying amplitudes. The stimulus waveform is shown at the top, and representative protrusion length changes are shown in the middle from a single vesicle where protrusion could be detected above the background created by probe binding to the pipette (A). Due to background fluorescence at the tip of the glass pipette, visualizing the protrusion length was not always possible. Shown below are the mean changes in vesicle fluorescence for W6NBD (B, n=3) and W7NBD (C, n=4). Responses to pressure stimuli exhibit a rapid phase that mirrors the stimulus, with an 80-ms rise time. This is followed by a slower exponential phase of gradual fluorescence increase. At the end of the stimulus, there were similar rapid and slow decay kinetics, with the fluorescence returning to background levels.

### Aspiration of cell-derived vesicles reveals inhomogeneity of tension reporting

We generated cell-derived vesicles from cultured cells and studied them in the presence of W7NBD. Supplementary Figure S6 illustrates an example of a pipette aspiration experiment with a vesicle from bovine aortic endothelial cells (BAEC). At 0 mmHg, the unperturbed vesicles exhibit barely detectable puncta, implying some membrane heterogeneity in the resting state. Upon applying suction to the pipette, the vesicles display more prominent puncta, albeit at different locations. This contrasts with synthetic GUVs, which exhibit homogeneous membrane labeling and tension sensitivity (Figure 8). Cell-derived vesicles (61) are a natural source of heterogeneous bilayers with domains that may affect the performance of the tension probe. This heterogeneity may represent domain structure where certain areas (i) have charge and lipid order characteristics that promote peptide binding, and (ii) can be unprotected and more prone to tension generation. The ability of W7NBD to highlight isolated domains warrants further investigation into the probe’s properties that can target specific domains.

## DISCUSSION

The developed probes, W6NBD and W7NBD, are based on a rigid scaffold of GsMTx4, the inhibitory cysteine knot peptide known to cause a rightward shift in the activation curves of channels responding to lipid tension (62). The peptide, stabilized by three internal disulfides, resembles a cone with a hydrophobic tip capped by two tryptophans (W6 and W7), which were chosen for insertion of NBD groups. Structurally similar to tryptophan, NBD fluorophores are more sensitive to the polarity of their environment. Based on the MD simulations (Figures 2 and 3), the rigid NBD-modified peptide maintains its shape but shifts its position and orientation within the bilayer, dynamically transitioning between a shallow adsorption state and a deeper submerged state. The occupancies of the two states depend on the lateral pressure of lipids, an inverse measure of membrane tension. When compared to the compressed state of the bilayer (50.7 Å²/lipid), the stretched state (65 Å²/lipid) alters the peptide’s orientation, pushes it deeper into the membrane, and reduces the likelihood of polar contacts for the NBD moiety by about twofold, based on all four simulated replicas of the peptide. The redistribution of the probe from the shallow to the deep state under tension increases the quantum yield, reflected by the overall fluorescence level. As shown in the timelines of parameters extracted from MD trajectories, the switching between deep and shallow states during bilayer compression or stretching is not an ‘all or none’ process; the probe is highly dynamic and occupies multiple intermediate states.

The experimental data presented here closely align with the computational predictions and provide strong, independent confirmation of the previously proposed mechanism of GsMTx4 action as a mobile amphipathic component that serves as a tension-dependent area tension-buffering reservoir to the exposed membrane leaflet (42). Because GsMTx4 is a modifier of native channel gating, its action is expected to occur in a physiological range of tensions.

At this stage, the behavior of the probes was tested primarily in model systems, including monolayers, small extruded liposomes, and giant unilamellar vesicles. Langmuir monolayer experiments well illustrated the probe action mechanism, albeit observed in reverse order. Monolayers allow for complete control of area and pressure, but lack the elastic coupling between inner and outer monolayers of a regular bilayer (63). Intercalated NBD peptides occupied a substantial area among the lipids but were pushed out within a narrow pressure range around the monolayer-bilayer equivalence point (35-40 mN/m). The slope of probe expulsion is directly related to the in-plane area occupied by the probe in the ‘deep’ state (Supplementary Figure S6), estimated at approximately 250 Å² (in the PC:PA:Chol mixture), which corresponds to about five lipid molecules. The rigid three-dimensional structure of the peptide determines this lipid-displacement area and the observed pressure (tension) sensitivity range, spanning 0-10 mN/m. The probe’s expulsion from the monolayer is reflected in a significant increase in compressibility and a concurrent decrease in fluorescence. Monolayer experiments also showed that probe incorporation and the sharpness of expulsion depend on lipid composition, especially the presence of cholesterol.

Electrostatic maps of simulated membranes with peptides consistently showed that local favorable interactions between multiple lysines and interfacial lipid groups can be offset by a large positive dipole potential within the membranes, thereby preventing deep penetration of a positively charged probe. The positive dipole potential appears to be the reason for the requirement of negatively charged lipids for both interfacial binding and deeper probe penetration. This is why the probes are completely indifferent to zwitterionic PC membranes.

SUVs containing a high mole fraction of anionic lipids, without ordering agents, immediately saturated the W6NBD fluorescence, indicating that, even without tension, the probe inserts in a deeply adsorbed state. The presence of lipid-ordering components such as sphingomyelin and especially cholesterol reduced the background insertion of the probes into liposomes and increased the dynamic range of responses to osmotically generated tension. We assume that the lipid-ordering effect of cholesterol works synergistically with its ability to increase the magnitude of the repulsive dipole potential (64).

Of the physiologically relevant lipids tested, PS and PA both act as efficient attractors to concentrate the probe on the membrane surface in a ‘shallow’ state and overcome the stiffness of ordering agents when tension was applied. PA was more effective at producing greater fluorescence change at lower concentrations (10-15 mole%) than PS, while still maintaining a low resting fluorescence in W6NBD. These two headgroups likely coordinate the peptide in different configurations due to their differing sizes, chemical compositions, distributions of ionizable groups, hydration levels, H-bonding capacities, and effective charges. When interacting with lysine residues, PA carries a -2e charge (65), and its cone shape (66) may cause curvature frustration that could also affect the depth of penetration. The lipid curvature-perturbing effects of shallowly adsorbed peptides have been analyzed in several studies (67–69). The prediction that GsMTx4 and its derivatives bind to the outer surface of the plasma membrane, imposing negative curvature, can be relevant to the curvature-sensing mechanism of Piezo channels, as these channels organize the membrane around themselves as positively invaginated pits (70).

The resting fluorescence levels and tension responses in SUVs were largely mirrored in aspiration experiments on GUVs. Timed osmotic dilution of SUVs using a stopped-flow machine enabled us to calibrate the fluorescence response of W6NBD based on estimated membrane tension. Liposomes containing phosphatidylserine and stabilized with sphingomyelin and cholesterol exhibited a 7% increase in fluorescence per mN/m during the initial phase of the dose-response curve. Similar liposomes containing phosphatidic acid, a more potent attractor, exhibited a 13% increase per mN/m. When tension was applied to GUVs, we observed a larger change per mN/m (15% in PS- and 33% in PA-containing membranes, respectively). The slope might be affected by differences between the two systems: in SUVs, we measure a population-averaged response that can be shallower due to slight population heterogeneity (the principle is discussed in (71)). In contrast, the optical response of a single uniform vesicle will not be smeared by the averaging effect. The slope of fluorescence versus tension was consistently higher when PA was present in the mixture than when PS was present, suggesting that, in the presence of PA, the effective displaced area (ΔA) associated with probe penetration is larger.

The time-resolved stopped-flow experiments also showed delayed fluorescence response kinetics (0.1-0.4 s), indicating the presence of slow stages, possibly involving diffusion-limited binding and transition to a deeply adsorbed state. The two-stage kinetics were clearly observed in giant vesicle (GUV) aspiration experiments: the fast stage followed the kinetics of tension onset (80 ms), likely reflecting the response of the pre-adsorbed probe, with the subsequent slow (∼0.5 s) stage likely representing the diffusion-limited arrival of the additional probe.

W7NBD appeared to be slightly more sensitive in both steady-state and dynamic tension responses in GUVs, although the overall response magnitude was smaller. This is likely due to the shallower resting depth of W7 (Supplemental Figure S2), which offers a different dynamic range for future studies. One key difference was that the resting depth of W7NBD appeared to be the same in both PS and PA containing vesicles, and this will be further examined through simulations and vesicle experiments.

The sensitivity of probe insertion to different electrostatic attractors and lipid-ordering components, such as cholesterol, clearly shows that, in real-world situations, the probe’s affinity must vary across different membrane domains, including liquid-disordered and liquid-ordered (raft) phases. Moreover, the data presented above demonstrate that the affinity for specific domains can change significantly with tension: the ordered phase might not accommodate the probe at rest, but under tension, that domain could become saturated with the probe. This is precisely what our first experiments with cell-derived vesicles suggest (Supplemental Figure S7). At resting (low) tension, the vesicle stains sparsely but unevenly, indicating the presence of more accessible domains. When tension is applied, a notable change in fluorescence is observed, localized in areas different from where the speckles were initially observed. This clearly indicates that the developed probes can be used to identify different domains and visualize tension distribution in a naturally nonuniform membrane.

## Materials and Methods

For full technical details, see Supplement, page 2.

### Analysis of GsMTx4 peptide, probe design, and simulations

Peptide amphipathicity was analyzed by mapping hydrophobicity and electrostatic potential on the solvent-accessible surface (Figure 1). Tryptophans W6 and W7, in salient positions crowning the hydrophobic end, were chosen as substitution sites for the NBD amino acid. The NBD moiety was first computationally incorporated in these two positions using CGenFF (72), and the peptide was then simulated in two lipid environments at different lateral pressures.

The lipids used to create membrane-forming mixtures for simulations and experiments included 1-palmitoyl-2-oleoyl-sn-glycero-3-phosphocholine (POPC), 1-palmitoyl-2-oleoyl-sn-glycero-3-phosphoglycerate (POPG), 1-palmitoyl-2-oleoyl-sn-glycero-3-phosphoserine (POPS), bovine heart Sphingomyelin (SM), and Cholesterol (Chol). The details of simulations are presented in the Supplemental document. The lipid bilayer was built of 128 lipids per leaflet, of two compositions, either POPC/POPG (1:1) or a mammalian-like mixture of POPC/POPS/Chol (6:2:2). A flexible orthogonal cell was simulated with periodic boundary conditions under 1 bar pressure and an initial lateral tension of 20 dyne/cm. All simulations were performed using NAMD2 (73) with the CHARMM27 force field (74), TIP3P water model, particle mesh Ewald method for long-range electrostatics estimation (75), a 10 Å cutoff for short-range electrostatic and Van der Waals forces, and a Langevin thermostat set at 310 K. All structural and statistical analyses for the MD simulations were performed using custom-written Tcl scripts in the VMD software. Electrostatic maps of membrane-embedded peptides were calculated using the PME modulus in VMD-NAMD.

### Peptide synthesis

The W6NBD and W7NBD GsMTx4 peptides were synthesized using solid-phase peptide synthesis (76), with the NBD amino acid (77) substituted for tryptophan. Commercial solid-state synthesis, followed by folding under natural oxidation of three internal disulfides, was performed by CSBio. The identity was verified with mass spectrometry.

### Fluorometric analysis of peptide response in osmotically shocked liposomes

Experiments were carried out on 0.2 µm extruded liposomes prepared from mixtures of POPC, POPG, POPS, POPA, sphingomyelin, and cholesterol (all from Avanti Polar Lipids) in specified molar ratios. To make liposomes osmotically sensitive, they were extruded in a 700 mOsm sorbitol buffer. Manual fluorometric analysis was conducted using a Horiba Fluoromax-4, with excitation set at 470 nm and emission at 554 nm. In stopped-flow experiments, osmotic shock fluorescence kinetics were recorded at 1 kHz over 20-second intervals using a BioLogic SFM2000 stopped-flow machine equipped with an 86 µL rectangular quartz cuvette. The illuminating beam entered the chamber at 90° relative to the PMT port from a 460 nm laser (MDL-III-460L, 30 mW, Opto Engine). The procedures are outlined in the text, legends to Figures 6 and 7, and in the Supplement.

### Monolayer fluorometry experiments

The Langmuir monolayer technique has been previously used to observe the behavior of unlabeled GsMTx4 and several of its mutants (42). A similar technique was used to study the behavior of the W6NBD and W7NBD fluorescent probes in a commercial Kibron mini-trough equipped with a laser and fluorometer, placed in a light-tight enclosure. For fluorometry, the same 460 nm laser was used as the excitation source. The emission light was filtered through a 525 nm long-pass filter (Edmund), collected using a lightguide-coupled collimating lens, and analyzed using a QEPRO-FL fluorometer (Ocean Optics). The background fluorescence measured before the experiment from the trough filled with pure subphase buffer was subtracted. The subphase used for all monolayer experiments was 10 mM phosphate buffer and 100 mM NaCl, pH 7.2. In all compression experiments, the W6NBD peptide was present in the subphase at a concentration of 0.3 µM. Lipids were deposited onto the subphase, and the solvent was allowed to evaporate for 30 minutes before compression was initiated. The intensity at the maximum emission wavelength of 448 nm was recorded and superimposed with the pressure isotherm on the area per lipid scale.

### Aspiration of giant unilamellar vesicles (GUVs)

Giant unilamellar vesicles (GUVs), 10-80 μm in diameter, were prepared by the electroformation technique as in (4). Once vesicles have formed, they are diluted with 0.75 ml of KHM imaging buffer (150 mM KCl, 20 mM Hepes/KOH pH 7.4, 1 mM MgCl2), mildly dispersed, and distributed to multiple viewing chambers. Chambers were placed on a Zeiss Axio Observer Z1 inverted microscope equipped with a Ludl motorized stage, Lumen Dynamics X-Cite fluorescence illuminator, and an Andor DU-897U-CSO back-illuminated electron multiplying camera. Vesicles were visualized with transmitted light through DIC optics and fluorescence with a 480/520 nm (ex/em) GFP filter set through a 63X 1.4 NA oil objective. The microscope and accessory equipment were controlled with Micromanager software. Fast kinetic imaging was achieved using QuBIO software to create the pressure steps and trigger the Andor camera acquisitions and the illuminator.

Pipettes were pulled and tips were broken to create openings with diameters of 5-10 μm. To prevent sticking, the tips were dipped in a solution of 95% dichlorobenzene and 5% dichlorodimethylsilane for 20 seconds, heated with a heat gun for an additional 20 seconds, and then dipped into a 1% solution of F-127 pluronic in sucrose-KHM (K+-HEPES-Mg Acetate) working solution for 2-3 minutes. Slight suction (0.5 mmHg) was used to keep the vesicle attached to the tip between suction steps. ALA clamp pressure was controlled by Micromanager software. The pipette and vesicle diameters, along with the protrusion length measured under specific applied pressure, were used to estimate membrane tension γ according to the equation from Rawicz (59).

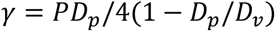

Where *P* is the pressure, *D_p_* is the pipette diameter, and *D_v_* is the radius of the vesicle.

### Cell membrane vesicles

Bovine aortic endothelial (BAEC) and C2C12 myoblast cells were grown to 80% confluence and treated with 25 mM formaldehyde and 2 mM DTT in normal saline (NaCl - 140 mM, HEPES - 10 mM, pH 7.4; KCl - 5 mM; CaCl2 – 2 mM; MgCl2 – 0.5 mM; glucose - 6 mM) for 2 hours at 37°C while gently shaking. Vesicles bud off and are suspended in the saline. 50 μL of vesicle saline was diluted into 1.5 mL of normal saline in preparation for aspiration experiments.

## Acknowledgment

This work was supported by NIH R21GM137274 grant to SS and TS.

U.S. Patent Application (USPTO No. 63/916,324) / Fluorescent Peptides For Membrane Tension Reporting / Suchyna et al. has been filed.

## Supplemental Material

## METHODS DETAILS

### Analysis of GsMTx4 peptide, probe design, and simulations

Peptide amphipathicity was analyzed by mapping hydrophobicity and electrostatic potential on the solvent-accessible surface. Tryptophans W6 and W7, in salient positions crowning the hydrophobic end, were chosen as substitution sites for the NBD dye. The NBD moiety was first computationally incorporated in these two positions using CGenFF (1), and the peptide was then simulated in two lipid environments at different lateral pressures.

The lipids used to create membrane-forming mixtures for simulations and experiments included 1-palmitoyl-2-oleoyl-sn-glycero-3-phosphocholine (POPC), 1-palmitoyl-2-oleoyl-sn-glycero-3-phosphoglycerate (POPG), 1-palmitoyl-2-oleoyl-sn-glycero-3-phosphoserine (POPS), bovine heart Sphingomyelin (SM), and Cholesterol (Chol). The details of simulations are presented the Supplemental document after following Figure S1. The lipid bilayer was built of 128 lipids per leaflet, of two compositions, either POPC/POPG (1:1) or a mammalian-like mixture of POPC/POPS/Chol (6:2:2). To increase the sample size, four W6NBD peptides were placed in the corners of the simulation cell, two on each side, on different diagonals. The membrane with peptides was hydrated, and Na^+^ and Cl^-^ - ions were added to create a 100 mM NaCl solution. Initial simulations were performed using the PC/PG system, but the production runs were conducted in the POPC/POPS/Chol mixture, which corresponds to the experimental conditions of monolayer and liposome experiments.

A flexible orthogonal cell was simulated with periodic boundary conditions under 1 bar pressure and an initial lateral tension of 20 dyne/cm. To mimic the compressed and stretched states of the bilayer, the simulation cell was initially contracted to approximately 45 Å^2^/lipid, causing the bilayer to start buckling, and then gradually expanded to roughly 68 Å^2^/lipid, where bilayer thinning was observed. Densities of 51 Å^2^/lipid and 65A^2^/lipid were selected as appropriate representations of a compressed (resting) and stretched state of the bilayer.

All simulations were performed using NAMD2 (2) with the CHARMM27 force field (3), TIP3P water model, particle mesh Ewald method for long-range electrostatics estimation (4), a 10 Å cutoff for short-range electrostatic and Van der Waals forces, and a Langevin thermostat set at 310 K. After the equilibration stage was completed, the system was simulated with fixed areas per lipid for 100 ns, and the trajectory was used to quantify the angular orientation of the peptide, statistics of contacts with water, and the distances of the peptide’s center of mass and the NBD group from the membrane midplane. Visualization of the peptides in the membrane was performed using Visual Molecular Dynamics (VMD) (5). Electrostatic maps of membrane-embedded peptides were calculated using the PME modulus in VMD-NAMD. All structural and statistical analyses for the MD simulations were performed using custom Tcl scripts in VMD.

### Peptide synthesis

The W6NBD and W7NBD GsMTx4 peptides were then synthesized using solid-phase peptide synthesis (6), with the NBD amino acid (7) substituted for tryptophan. Commercial solid-state synthesis, followed by folding under natural oxidation of three internal disulfides, was performed by CSBio. The identity was verified with mass spectrometry.

### Fluorometric analysis of peptide response in osmotically shocked liposomes

Lipid mixtures were prepared from chloroform solutions of POPC, POPG, POPS, POPA, sphingomyelin, and cholesterol (all from Avanti Polar Lipids) in specified molar ratios and measured out to a final weight of 5 mg. The mixtures were then dried using N_2_ gas and desiccated overnight in a vacuum. The next day, lipid films were suspended in 1 mL of 0.5 M sorbitol, 100 mM NaCl, and 10 mM NaPi buffer (700 mOsm, pH 7.2), rehydrated for 30 minutes, and then extruded through a 0.2 µm nucleopore filter to create uniform unilamellar vesicles (SUVs). SUV size was confirmed using a DLS particle size analyzer (90Plus, Brookhaven Instruments). Freshly prepared liposomes were used within 2 days.

Manual fluorometric analysis was conducted using a Horiba Fluoromax-4, with excitation set at 470 nm and emission at 554 nm. A 30 µM stock solution of W6NBD GsMTx4 in the 700 mOsm sorbitol buffer was prepared and stored in 1 mL aliquots. Liposome samples and peptides were loaded into 4 mL (10×10 mm) quartz cuvettes. To maintain consistency with the monolayer experiments (below), 20 µL of W6NBD and 20 µL of liposome solution were added to 1.98 mL of sorbitol buffer, resulting in a final volume of 2 mL. First, baseline fluorescence was measured using a sample of pure W6NBD in sorbitol buffer and recorded for 90 seconds. Next, 20 µL of SUVs were added to evaluate the increase in fluorescence caused by peptide binding and insertion into membranes. In a subsequent experiment, liposomes and peptides were pre-mixed, added to the cuvette prior to measurement, and recorded for 60 seconds. This was followed by osmotic shock or dilutions, achieved by adding 200 µL, 500 µL, or 1 mL of distilled water, with another 60 seconds of recording. As a control for non-shock dilution, the same amount of 700 mOsm sorbitol buffer was added; fluorescence levels were measured, and the ratio of shock to non-shock fluorescence levels was calculated. The pre-shock level of fluorescence (counts per second) was multiplied by this ratio to quantify the effect of osmotic stress (see legend to Figure 5). In the bar graphs, all data are normalized to the level of fluorescence of pure peptide.

In stopped-flow experiments, osmotic shock fluorescence kinetics were recorded at 1 kHz over 20-second intervals using a BioLogic SFM2000 stopped-flow machine equipped with an 86 µL rectangular quartz cuvette. The illuminating beam entered the chamber at 90° relative to the PMT port. A 460 nm laser (MDL-III-460L, 30 mW, Opto Engine), coupled through a quartz light guide, served as the excitation light source. The excitation and emission paths were additionally filtered with 475 nm short-pass and 525 nm long-pass filters (Edmund Scientific), respectively. Syringe 1 contained liposomes, W6NBD (0.3 μM), in the sorbitol buffer used in the previous fluorometric analysis. Syringe 2 contained distilled water or sorbitol buffer for comparison of shock and dilutions. The total mixing volume was 302 μL, and the mixing time was 8 ms. Ratios of mixing volumes for syringe 1 to syringe 2 were as follows: 1:3, 1:2, 1:1, 2:1, 3:1, 4:1, 5:1, and 6:1. The background, recorded from pure buffer with NBD (∼5% of the signal), was subtracted, and the fluorescence traces were normalized at appropriate dilution ratios to the unshocked controls, where liposomes were mixed with the sorbitol buffer in which they were extruded.

### Monolayer fluorometry experiments

The Langmuir monolayer technique has been previously used to observe the behavior of unlabeled GsMTx4 and several of its mutants (8). A similar technique was used to study the behavior of W6NBD and W7NBD fluorescent probes in a commercial Kibron mini-trough, equipped with a laser, fluorometer, and placed in a light-tight enclosure. For fluorometry, the same 460 nm laser was used as a source of the excitation light. The end of the light guide features a lens that defocuses the beam to a spot of 25 mm in diameter, along with a short-pass 475 nm filter that blocks the spill into the emission range. The emission light was filtered through a 525 nm long-pass filter (Edmund), collected using a lightguide-coupled collimating lens, and analyzed using a QEPRO-FL fluorometer (Ocean Optics). The background fluorescence measured before the experiment from the trough filled with pure subphase buffer was subtracted. The subphase used for all monolayer experiments was 10 mM phosphate buffer and 100 mM NaCl, pH 7.2. In all compression experiments, the W6NBD peptide was present in the subphase at a concentration of 0.3 µM. Lipid mixtures listed in figures were diluted in 9:1 chloroform:methanol solutions to a working concentration of 0.5 mg/mL. Lipids were deposited onto the subphase, and the solvent was allowed to evaporate for 10 minutes before compression was initiated. For experiments with W6NBD in the subphase, this evaporation time period was extended to 30 minutes to allow the peptide to equilibrate with the monolayer. Compressions were performed up to the point of collapse over 7-minute periods. The intensity at the maximum emission wavelength of 448 nm was recorded and superimposed with the pressure isotherm on the area per lipid scale.

### Aspiration of giant unilamellar vesicles (GUVs)

Giant unilamellar vesicles (GUVs), 10-80 μm in diameter, were prepared by the electroformation technique as in (4). Chloroform stocks of POPC, POPG, and cholesterol were 25 mg/ml, and POPS, PA, and Porcine Brain Sphingomyelin (BSM) were 10 mg/ml (all from Avanti Polar Lipids). Liposome compositions were calculated using the Liposome Calculator App developed by Roman Volinsky. Platinum wires were coated with lipid solutions pre-mixed in specific ratios, dried for 1 hr under vacuum, and placed in chambers with 0.5 ml of 200 mM sucrose. Electroformation occurred under a 2.2-volt p-p 10 Hz sine wave for 1 hr, followed by a 2.4-volt p-p sine wave at 2 Hz for an additional 30 min. Coverslips in imaging chambers were precoated with BSA for 1 hr, rinsed with water, and air-dried. Once vesicles have formed, they are diluted with 0.75 ml of KHM imaging buffer (150 mM KCl, 20 mM Hepes/KOH pH 7.4, 1 mM MgCl2), mildly dispersed, and distributed to multiple viewing chambers. Chambers were placed on a Zeiss Axio Observer Z1 inverted microscope equipped with a Ludl motorized stage, Lumen Dynamics X-Cite fluorescence illuminator, and an Andor DU-897U-CSO back-illuminated electron multiplying camera. Vesicles were visualized with transmitted light through DIC optics and fluorescence with a 480/520 nm (ex/em) GFP filter set through a 63X 1.4 NA oil objective. The microscope and accessory equipment were controlled with Micromanager software. Fast kinetic imaging was achieved using QuBIO software to create the pressure steps and trigger the Andor camera acquisitions and the illuminator.

Pipettes were pulled using a Narishige NP-7 horizontal puller, and tips were broken to create openings with diameters of 5-10 μm. Tips were heat-polished and bent to a 15° angle so that, on the microscope stage holder, they would be nearly parallel to the surface of the cover slip. To prevent sticking, the tips were dipped in a solution of 95% dichlorobenzene and 5% dichlorodimethylsilane for 20 seconds, heated with a heat gun for an additional 20 seconds, and then dipped into a 1% solution of F-127 pluronic in sucrose-KHM (K+-HEPES-Mg Acetate) working solution for 2-3 minutes. All pluronic solution was replaced with a 1% BSA sucrose-KHM solution by using a syringe to force it from the tip before use. Pipettes were mounted on a holder on a Sutter micromanipulator with an ALA pressure clamp port for applying suction.

Slight suction (0.5 mmHg) was used to keep the vesicle attached to the tip between suction steps. ALA clamp pressure was controlled by Micromanager software. The pipette and vesicle diameters, along with the protrusion length measured under specific applied pressure, were used to estimate membrane tension γ according to the equation from Rawicz (9).

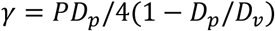

Where *P* is the pressure, *D_p_* is the pipette diameter, and *D_v_* is the radius of the vesicle.

---

**Figure S1.**
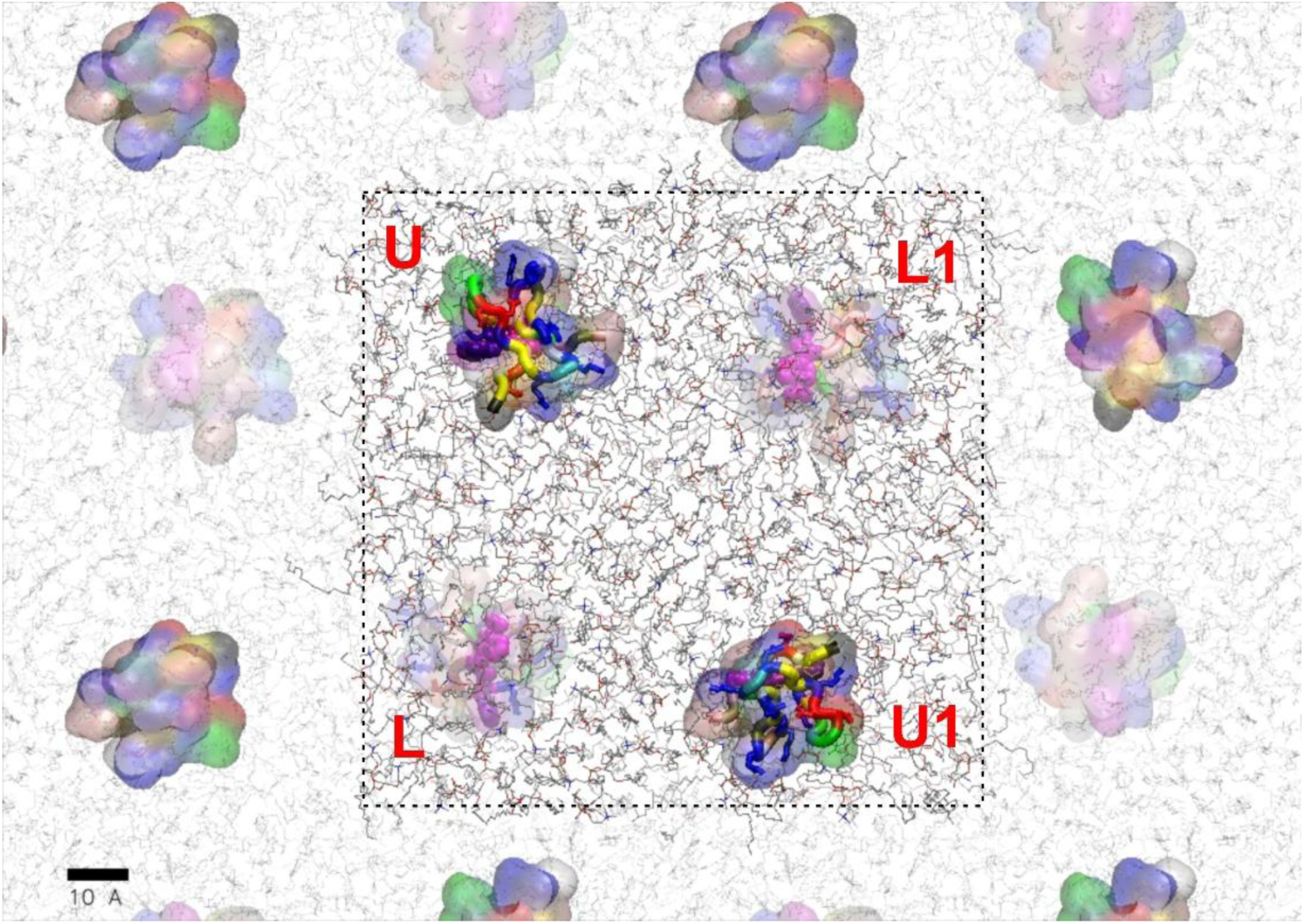
Arrangement of the simulation cell containing a rectangular membrane slab (128 lipids per side) and four copies of W6NBD peptide (U, U1, L, L1) placed at two opposite corners on each side. The periodic images of the simulation cell are also shown. Following pilot runs, the area of the lipid slab was set to satisfy 50.7 Å²/lipid for the compressed and 65 Å²/lipid for the expanded state. The system was allowed to equilibrate at each lateral dimension for 100 ns, after which a 100 ns production run was completed. Snapshots of typical positions of W6NBD in compressed and expanded bilayers are shown in Figure 2 of the main text.

**Figure S2.**
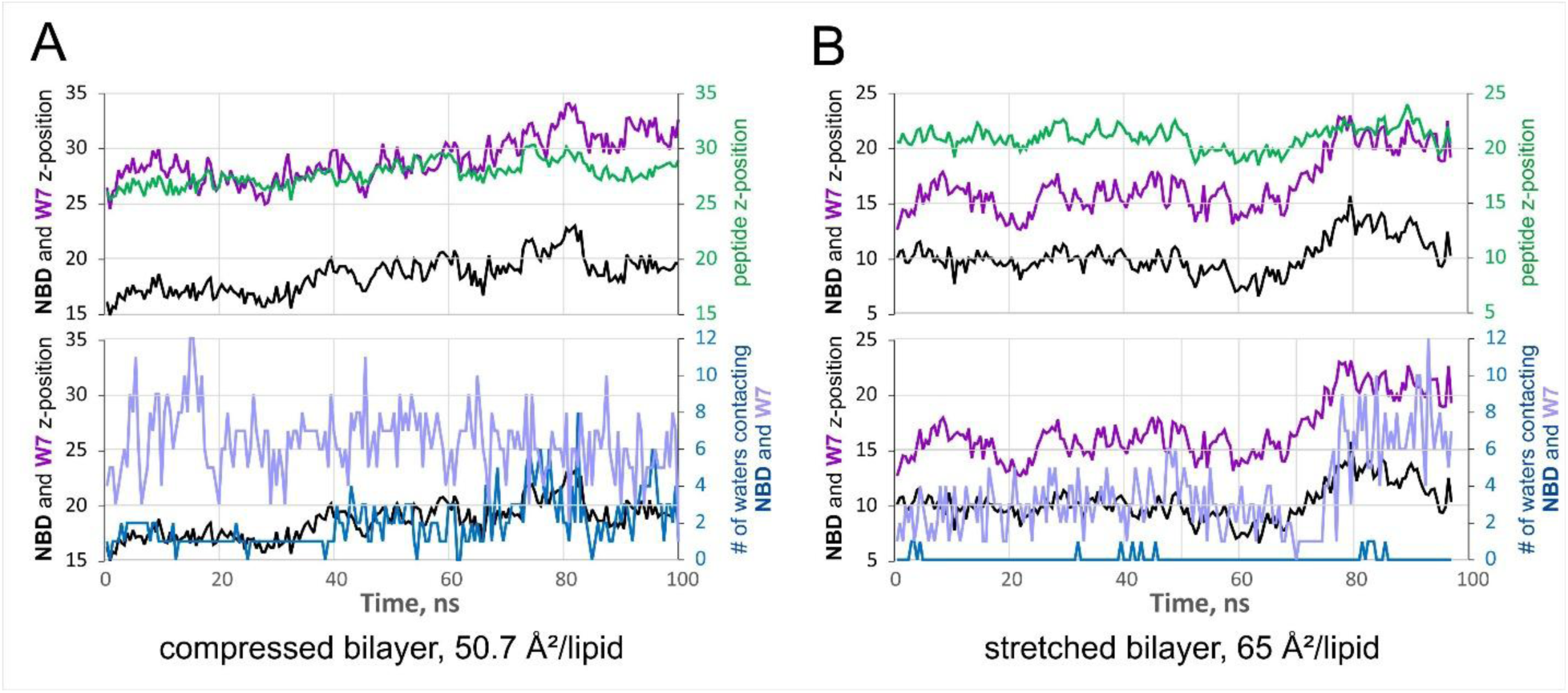
Positions of W7 sidechains in W6NBD simulations in the compressed (A) and stretched (B) bilayer. The graphs show the time courses of the same parameters as in Fig. 3 (main text), with the positions of the sidechain center of mass of W7 (purple traces) relative to the center of mass of NBD6 (black traces). The center of mass position for the entire peptide is represented by the green line, and the numbers of water molecules surrounding the W7 and NBD6 groups are depicted by the violet and blue traces, respectively.

**Table S1.** The summary of parameters related to the conformation of W6NBD. The parameters were extracted from a 100-ns simulation of a simulation cell with four replicas of W6NBD peptides (Figure S1). The parameters are compared for the W6NBD group with the W7 side chain present in the same peptide, in the same simulation. The comparison of the upper and lower halves of the table shows that W6NBD is immersed deeper and has substantially fewer contacts with water, especially in the stretched bilayer at 65 Å² per lipid.

| Residue | Area per lipid, Å<sup>2</sup> | replica name | peptide z-position | # of contacting waters | NBD z-position | peptide angle |
| --- | --- | --- | --- | --- | --- | --- |
| W6NBD | 50.7 | U | 24.8 | 0.51 | 14.8 | 37.3 |
|  |  | U1 | 26.2 | 0.80 | 16.1 | 36.6 |
|  |  | L | 25.1 | 0.09 | 14.4 | 49.3 |
|  |  | L1 | 27.7 | 1.94 | 18.5 | 67.5 |
|  |  | Average | 26.0 | 0.83 | 16.0 | 47.7 |
|  | 65.0 | U | 20.8 | 0.12 | 10.8 | 40.4 |
|  |  | U1 | 20.2 | 0.51 | 10.4 | 41.7 |
|  |  | L | 21.0 | 0.07 | 10.2 | 42.2 |
|  |  | L1 | 22.6 | 0.96 | 13.3 | 51.8 |
|  |  | Average | 21.1 | 0.42 | 11.2 | 44.0 |
|  | 65.0 - 50.7 | Average | -19% | -50% | -30% | -8% |
| W7 | 50.7 | U | 24.8 | 2.54 | 20.0 | 53.9 |
|  |  | U1 | 26.2 | 5.38 | 21.5 | 54.2 |
|  |  | L | 25.1 | 6.18 | 22.3 | 66.2 |
|  |  | L1 | 27.7 | 6.37 | 29.1 | 86.6 |
|  |  | Average | 26.0 | 5.12 | 23.2 | 65.2 |
|  | 65.0 | U | 20.8 | 2.49 | 15.9 | 54.9 |
|  |  | U1 | 20.2 | 3.47 | 15.9 | 58.0 |
|  |  | L | 21.0 | 3.65 | 16.7 | 58.3 |
|  |  | L1 | 22.6 | 3.88 | 18.2 | 62.8 |
|  |  | Average | 21.1 | 3.38 | 16.7 | 58.5 |
|  | 65.0 - 50.7 | Average | -19% | -34% | -28% | -10% |

**Figure S3.**
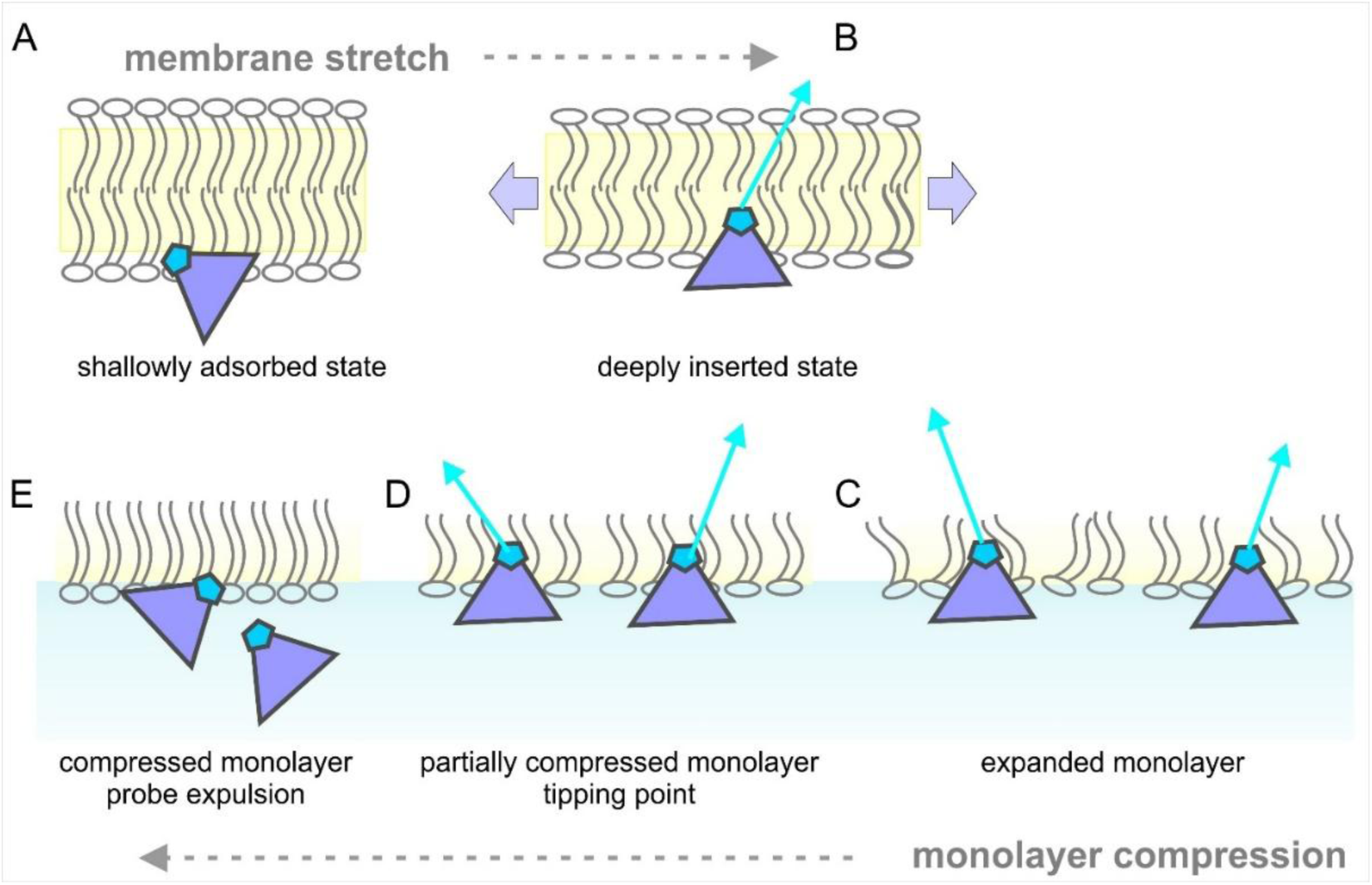
The relationship between the processes observed in monolayer compression and membrane stretching experiments. (A) In a resting, compressed membrane, the probe remains in a shallow absorbed state, resulting in low fluorescence. When the membrane is stretched (B), the probe penetrates deeper, causing an increase in fluorescence. Compression experiments (C-E) always start with an expanded monolayer (C), which allows the probe to be in a deeply inserted position. Compression to about 30 mN/m condenses the monolayer, reaching maximum stiffness and fluorescence at the tipping point (D). Beyond this point, expulsion begins, marked by a decrease in fluorescence and an increase in compressibility (E). The monolayer experiment is analogous to probe expulsion from a stretched membrane when tension is released, because lateral pressure is the inverse of membrane tension.

**Figure S4.**
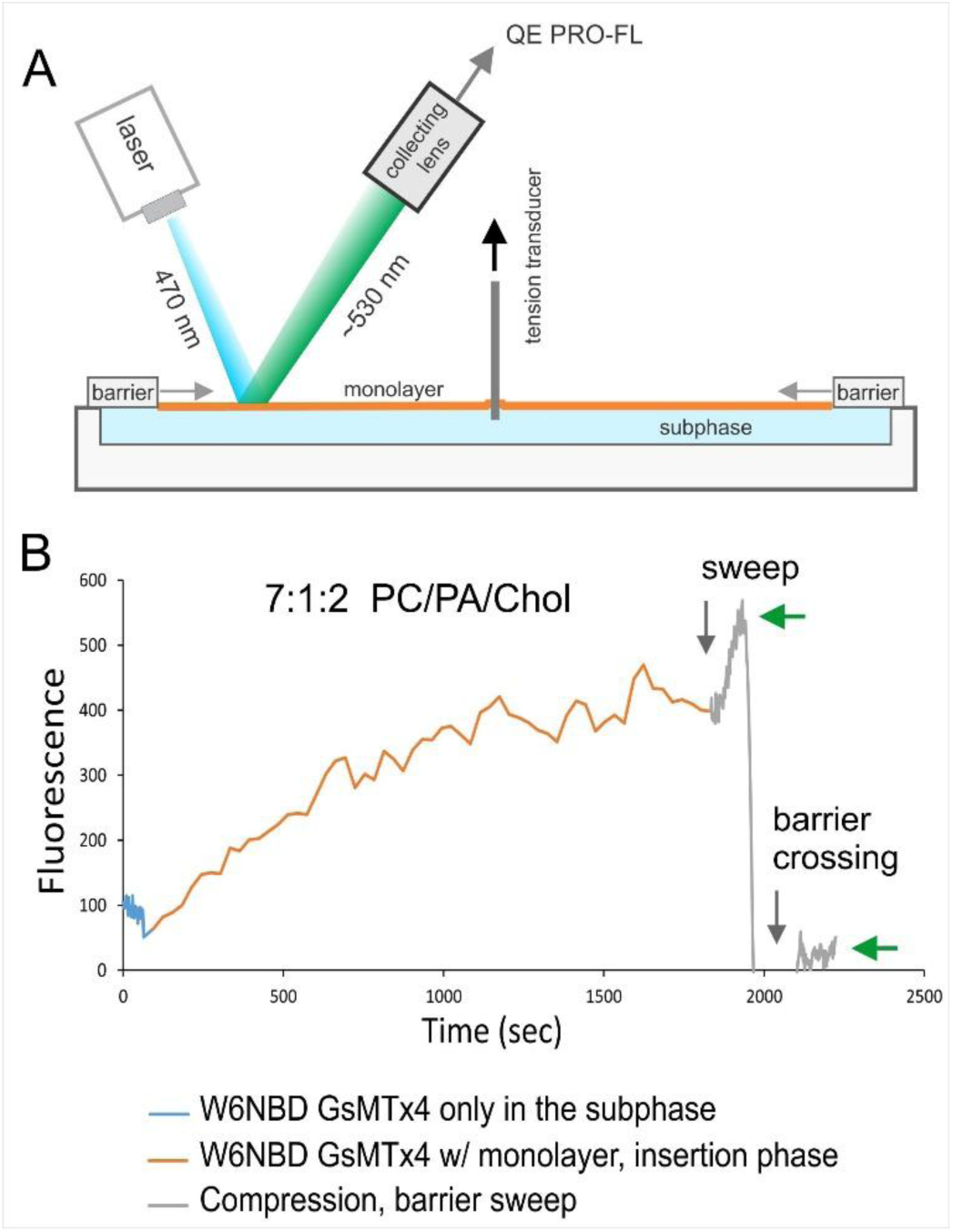
Measurement of W6NBD incorporation into the lipid monolayer. The ‘monolayer sweep’ experiment determined the extent to which the probe incorporates into a relaxed monolayer and the amount that remains in the subphase. A commercial monolayer setup (Kibron Microtrough) was adapted for optical measurements by incorporating a 470 nm laser and a QE PRO-FL spectrophotometer equipped with a light guide and a collecting lens (A). The laser beam was focused at a spot about 4 cm from the edge of the monolayer trough, illuminating the area that will be passed by the barrier during compression. The W6NBD probe was dissolved in the subphase buffer at 0.3 µM, and the aqueous probe fluorescence was measured before lipid application. The amount of lipid was sufficient to create an expanded monolayer at a surface pressure of ∼10 mN/m. The kinetics of probe incorporation into this monolayer over 30 min are shown in panel B (orange trace). Then, the barriers started moving (as indicated by a gray vertical arrow), causing a compression that visibly increased fluorescence due to the compaction of the lipid with the probe (gray stretch). When the barrier crossed the beam, the signal dropped to zero (blackout), and when the barrier ultimately passed the beam, sweeping the lipids to the center, the level of fluorescence dropped to ∼5% of the value before the sweep (green arrows that designate the fluorescence before and after the blackout due to barrier crossing the beam). This result illustrates that 95% of the probe got incorporated into the lipid film, albeit with relatively slow kinetics.

**Figure S5.**
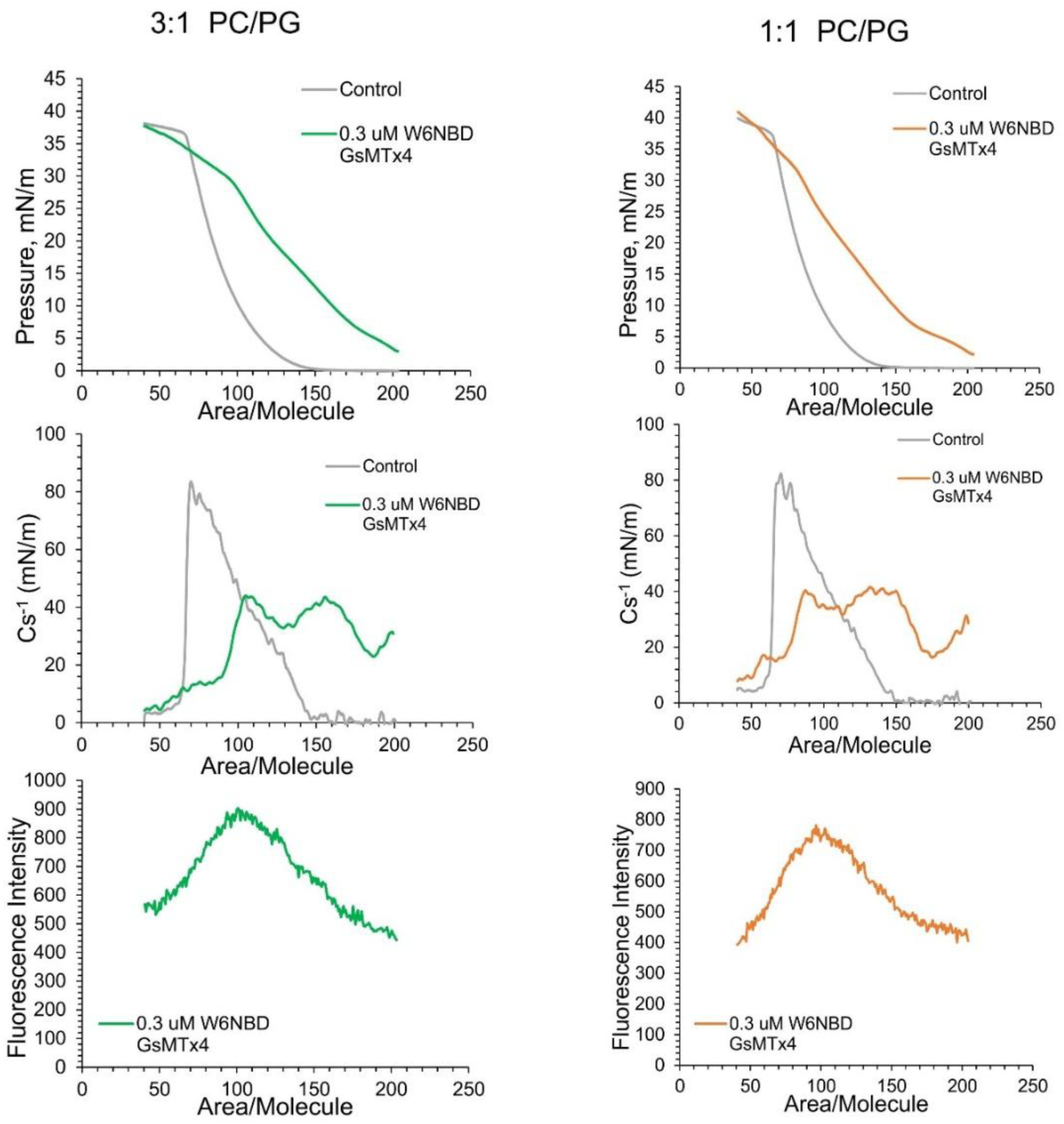
Compression isotherms for monolayers formed of binary PC/PG (3:1, panel A) and (1:1, panel B) mixtures with incorporated W6NBD. Gray traces represent controls without the peptide; blue traces represent experiments with 3×10^-7^ M W6NBD in the subphase. After spreading the lipid, the fluorescent peptide was allowed to incorporate for 30 minutes. The upper row represents the original compression isotherms, which visibly bend in the presence of W6NBD at pressures between 30 and 35 mN/m. Peaks in the plots of inverted compression modulus (Cs^-1^, second row) indicate this irregularity in the slope. The drop of Cs^-1^ at this point reflects an increase in compressibility related to the beginning of the expulsion of the peptide from the monolayer. This expulsion is confirmed by the simultaneous drop in fluorescence (bottom row) as the compression proceeds.

**Figure S6.**
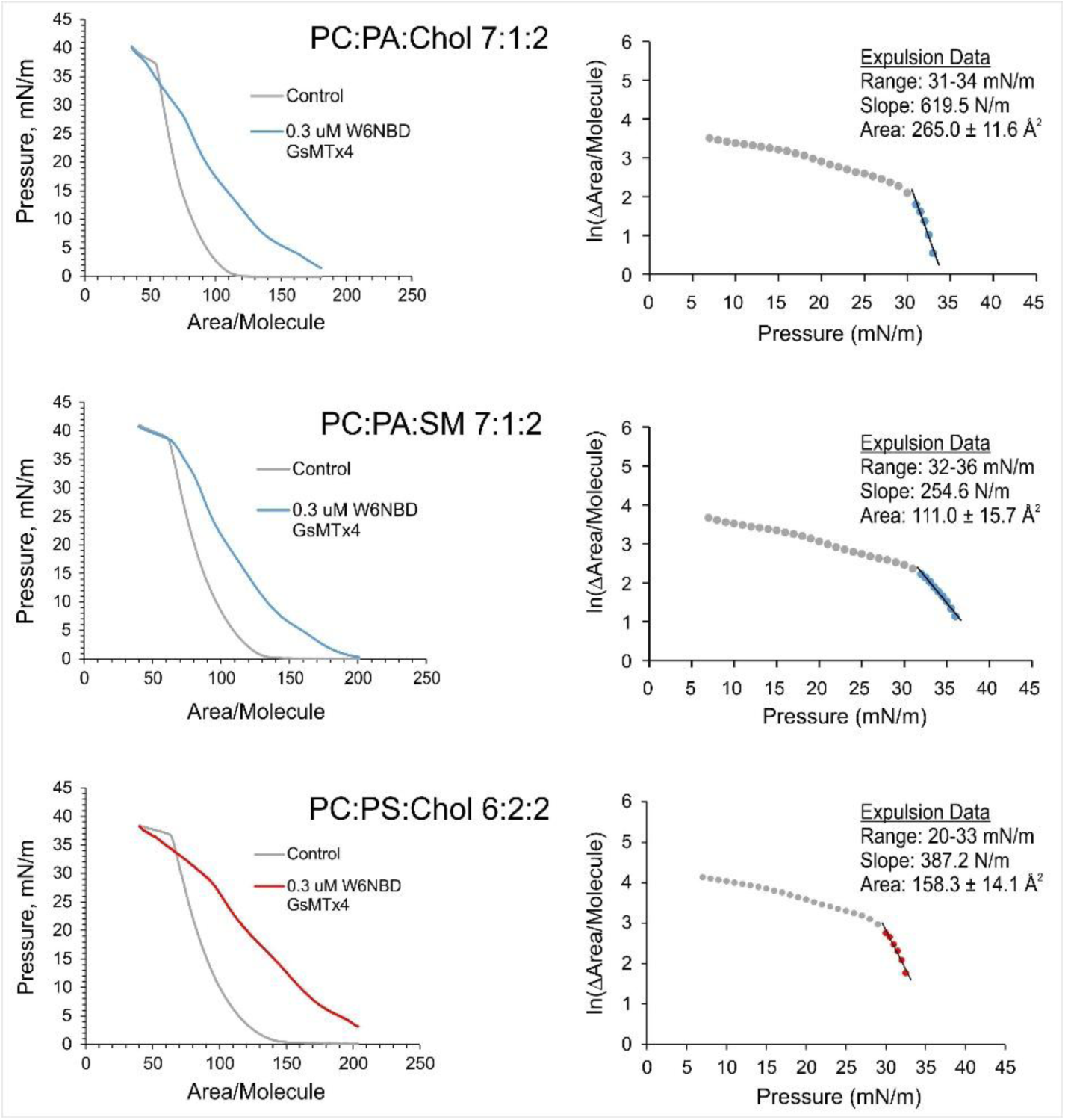
Estimation of the insertion/expulsion molecular area of W6NBD in monolayers of three different compositions. The excess area of the monolayer with the ‘guest’ peptide reflects the mole fraction of the ‘guest’, and the slope of the monolayer area decrease with lateral pressure directly indicates the mechanical component of the equilibrium and the area taken by individual ‘guest’ molecules in the plane of the monolayer. One can write a Boltzmann-type equation for the equilibrium constant between a deeply inserted state with probability (*p_lip_*) and an expelled state with probability (*p_bulk_*). *ΔG* is the chemical energy separating the monolayer-inserted and soluble states, and *πa* is the pressure-dependent component of the free energy difference.

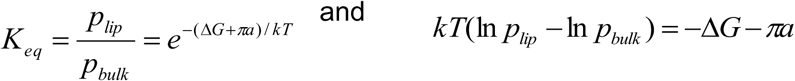 Differentiating with respect to *π,* we obtain

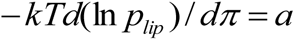 The area determination procedure involves re-plotting the pressure-area isotherms in area-pressure coordinates, finding the difference in area as a function of pressure, taking the log of ΔA, and fitting the slope. The slope and deduced areas depend on the presence of lipid-ordering components, with the lowest (∼110 Å²) observed in the PC:PA:SM mixture and the highest (∼250 Å²) recorded in the presence of Cholesterol. The data are based on 3 independent experiments.

**Figure S7.**
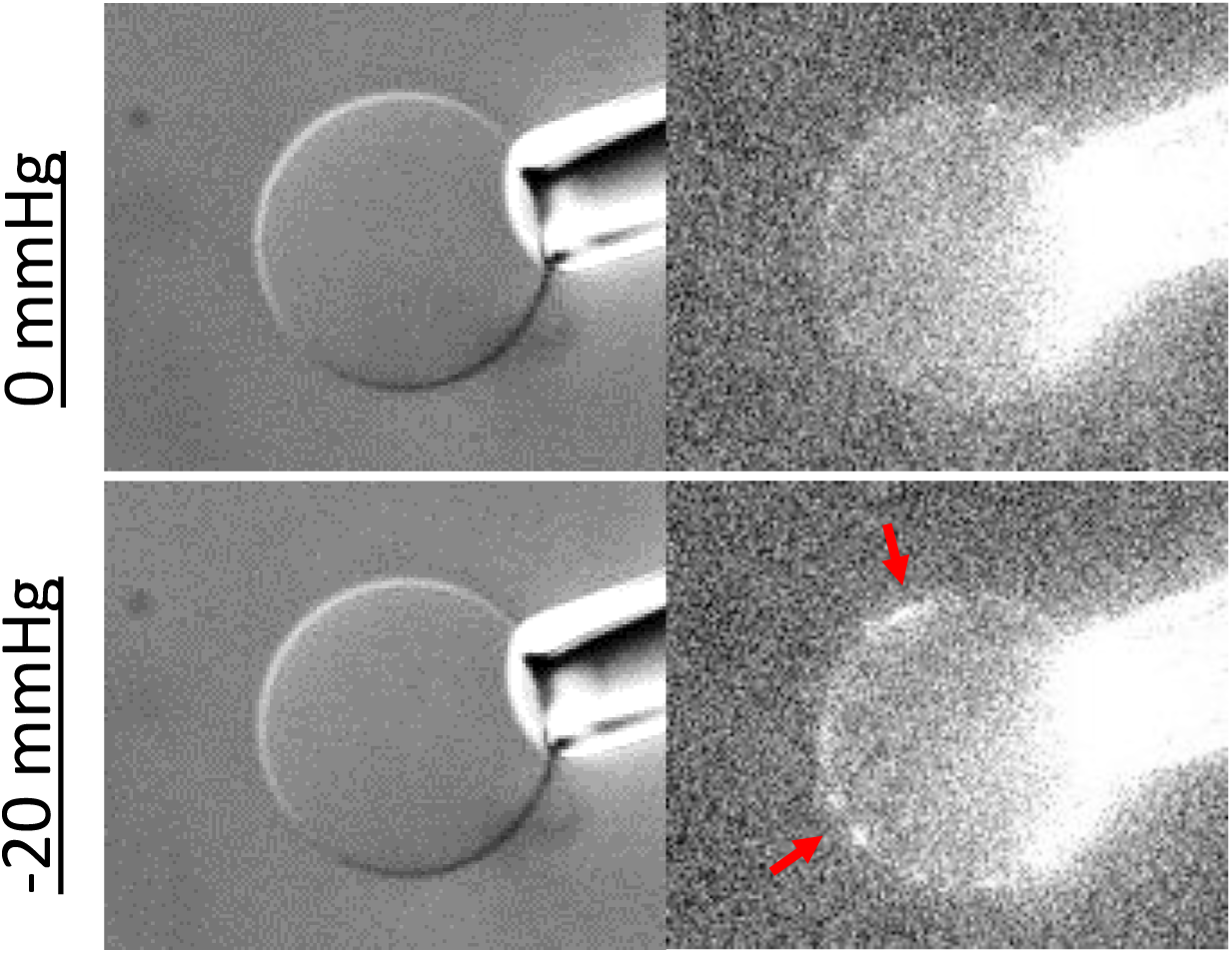
Pipette-aspiration experiment with vesicles derived from bovine endothelial BAEC cells. The vesicles were generated using formaldehyde and DTT treatment. The images were acquired in the presence of 0.4 μM W7NBD.

## References

1. F. Meng, F. Sachs, Visualizing dynamic cytoplasmic forces with a compliance-matched FRET sensor. J Cell Sci 124, 261–269 (2011).

2. F. Meng, T. M. Suchyna, F. Sachs, A fluorescence energy transfer-based mechanical stress sensor for specific proteins in situ. FEBS J. 275, 3072–3087 (2008).

3. C. Grashoff et al., Measuring mechanical tension across vinculin reveals regulation of focal adhesion dynamics. Nature 466, 263–266 (2010).

4. X. Wang, T. Ha, Defining single molecular forces required to activate integrin and notch signaling. Science 340, 991–994 (2013).

5. A. L. Cost, P. Ringer, A. Chrostek-Grashoff, C. Grashoff, How to Measure Molecular Forces in Cells: A Guide to Evaluating Genetically-Encoded FRET-Based Tension Sensors. Cell Mol Bioeng 8, 96–105 (2015).

6. T. R. Ham, K. L. Collins, B. D. Hoffman, Molecular Tension Sensors: Moving Beyond Force. Curr Opin Biomed Eng 12, 83–94 (2019).

7. E. R. Rojas, K. C. Huang, Regulation of microbial growth by turgor pressure. Curr Opin Microbiol 42, 62–70 (2018).

8. E. Atilgan, V. Magidson, A. Khodjakov, F. Chang, Morphogenesis of the Fission Yeast Cell through Cell Wall Expansion. Curr Biol 25, 2150–2157 (2015).

9. W. Fricke, M. C. Jarvis, C. T. Brett, Turgor pressure, membrane tension and the control of exocytosis in higher plants. Plant Cell Environ 23, 999–1003 (2000).

10. N. C. Gauthier, T. A. Masters, M. P. Sheetz, Mechanical feedback between membrane tension and dynamics. Trends Cell Biol 22, 527–535 (2012).

11. P. Nassoy, C. Lamaze, Stressing caveolae new role in cell mechanics. Trends Cell Biol 22, 381–389 (2012).

12. J. Dai, M. P. Sheetz, Axon membrane flows from the growth cone to the cell body. Cell 83, 693–701 (1995).

13. F. M. Hochmuth, J. Y. Shao, J. Dai, M. P. Sheetz, Deformation and flow of membrane into tethers extracted from neuronal growth cones. Biophys J 70, 358–369 (1996).

14. A. D. Lieber, S. Yehudai-Resheff, E. L. Barnhart, J. A. Theriot, K. Keren, Membrane tension in rapidly moving cells is determined by cytoskeletal forces. Curr Biol 23, 1409–1417 (2013).

15. Z. Shi, Z. T. Graber, T. Baumgart, H. A. Stone, A. E. Cohen, Cell Membranes Resist Flow. Cell 175, 1769–1779 e1713 (2018).

16. R. Dharan et al., Intracellular pressure controls the propagation of tension in crumpled cell membranes. Nature communications 16, 91 (2025).

17. D. P. Corey, N. Akyuz, J. R. Holt, Function and Dysfunction of TMC Channels in Inner Ear Hair Cells. Cold Spring Harb Perspect Med 9 (2019).

18. C. A. Haselwandter, R. MacKinnon, Piezo’s membrane footprint and its contribution to mechanosensitivity. Elife 7 (2018).

19. C. D. Cox, N. Bavi, B. Martinac, Origin of the Force: The Force-From-Lipids Principle Applied to Piezo Channels. Curr Top Membr 79, 59–96 (2017).

20. J. M. Kefauver, A. B. Ward, A. Patapoutian, Discoveries in structure and physiology of mechanically activated ion channels. Nature 587, 567–576 (2020).

21. R. Syeda et al., Piezo1 Channels Are Inherently Mechanosensitive. Cell reports 17, 1739–1746 (2016).

22. S. G. Brohawn, Z. Su, R. MacKinnon, Mechanosensitivity is mediated directly by the lipid membrane in TRAAK and TREK1 K+ channels. Proc Natl Acad Sci U S A 111, 3614–3619 (2014).

23. S. H. Woo, E. A. Lumpkin, A. Patapoutian, Merkel cells and neurons keep in touch. Trends Cell Biol 25, 74–81 (2015).

24. A. H. Lewis, M. E. Cronin, J. Grandl, Piezo1 ion channels are capable of conformational signaling. Neuron 112, 3161–3175 e3165 (2024).

25. S. S. Ranade et al., Piezo1, a mechanically activated ion channel, is required for vascular development in mice. Proc Natl Acad Sci U S A 111, 10347–10352 (2014).

26. B. Xiao, Levering Mechanically Activated Piezo Channels for Potential Pharmacological Intervention. Annu Rev Pharmacol Toxicol 60, 195–218 (2020).

27. A. H. Lewis, J. Grandl, Mechanical sensitivity of Piezo1 ion channels can be tuned by cellular membrane tension. Elife 4 (2015).

28. C. D. Cox et al., Removal of the mechanoprotective influence of the cytoskeleton reveals PIEZO1 is gated by bilayer tension. Nature communications 7, 10366 (2016).

29. B. Sorum, T. Docter, V. Panico, R. A. Rietmeijer, S. G. Brohawn, Tension activation of mechanosensitive two-pore domain K+ channels TRAAK, TREK-1, and TREK-2. Nature communications 15, 3142 (2024).

30. J. R. Holt et al., Spatiotemporal dynamics of PIEZO1 localization controls keratinocyte migration during wound healing. Elife 10 (2021).

31. M. M. Kozlov, L. V. Chernomordik, Membrane tension and membrane fusion. Curr Opin Struct Biol 33, 61–67 (2015).

32. T. Parasassi, E. Gratton, W. M. Yu, P. Wilson, M. Levi, Two-photon fluorescence microscopy of laurdan generalized polarization domains in model and natural membranes. Biophys J 72, 2413–2429 (1997).

33. E. K. Krasnowska, E. Gratton, T. Parasassi, Prodan as a membrane surface fluorescence probe: partitioning between water and phospholipid phases. Biophys J 74, 1984–1993 (1998).

34. T. Parasassi, E. K. Krasnowska, L. Bagatolli, E. Gratton, LAURDAN and PRODAN as polarity-sensitive fluorescent membrane probes. J Fluoresc 8, 365–373 (1998).

35. Y. L. Zhang, J. A. Frangos, M. Chachisvilis, Laurdan fluorescence senses mechanical strain in the lipid bilayer membrane. Biochem Biophys Res Commun 347, 838–841 (2006).

36. M. A. Boyd, N. P. Kamat, Visualizing Tension and Growth in Model Membranes Using Optical Dyes. Biophys J 115, 1307–1315 (2018).

37. E. Zapata-Mercado, E. V. Azarova, K. Hristova, Effect of osmotic stress on live cell plasma membranes, probed via Laurdan general polarization measurements. Biophys J 121, 2411–2418 (2022).

38. A. Colom et al., A fluorescent membrane tension probe. Nat Chem 10, 1118–1125 (2018).

39. C. Roffay et al., Tutorial: fluorescence lifetime microscopy of membrane mechanosensitive Flipper probes. Nat Protoc 19, 3457–3469 (2024).

40. T. M. Suchyna, Piezo channels and GsMTx4: Two milestones in our understanding of excitatory mechanosensitive channels and their role in pathology. Prog Biophys Mol Biol 10.1016/j.pbiomolbio.2017.07.011 (2017).

41. C. Bae, F. Sachs, P. A. Gottlieb, The mechanosensitive ion channel Piezo1 is inhibited by the peptide GsMTx4. Biochemistry 50, 6295–6300 (2011).

42. R. Gnanasambandam et al., GsMTx4: Mechanism of Inhibiting Mechanosensitive Ion Channels. Biophys J 112, 31–45 (2017).

43. S. Feryforgues, J. P. Fayet, A. Lopez, Drastic Changes in the Fluorescence Properties of Nbd Probes with the Polarity of the Medium - Involvement of a Tict State. J Photoch Photobio A 70, 229–243 (1993).

44. R. E. Oswald, T. M. Suchyna, R. McFeeters, P. Gottlieb, F. Sachs, Solution structure of peptide toxins that block mechanosensitive ion channels. J Biol Chem 277, 34443–34450 (2002).

45. K. Nishizawa et al., Effects of Lys to Glu mutations in GsMTx4 on membrane binding, peptide orientation, and self-association propensity, as analyzed by molecular dynamics simulations. Biochim Biophys Acta 1848, 2767–2778 (2015).

46. W. C. Wimley, S. H. White, Experimentally determined hydrophobicity scale for proteins at membrane interfaces. Nat.Struct.Biol. 3, 842–848 (1996).

47. A. S. Ladokhin, S. Jayasinghe, S. H. White, How to measure and analyze tryptophan fluorescence in membranes properly, and why bother? Anal Biochem 285, 235–245 (2000).

48. F. Sun, W. Zong, R. Liu, J. Chai, Y. Liu, Micro-environmental influences on the fluorescence of tryptophan. Spectrochim Acta A Mol Biomol Spectrosc 76, 142–145 (2010).

49. S. C. Haldar, A., “Application of NBD-labeled lipids in membrane and cell biology” in Fluorescent Methods to Study Biological Membranes. (Springer, 2012), pp. 37–50.

50. H. Brockman, Dipole potential of lipid membranes. Chem Phys Lipids 73, 57–79 (1994).

51. F. Zhou, K. Schulten, Molecular-Dynamics Study of A Membrane Water Interface. Journal of Physical Chemistry 99, 2194–2207 (1995).

52. K. Gawrisch et al., Membrane dipole potentials, hydration forces, and the ordering of water at membrane surfaces. Biophys J 61, 1213–1223 (1992).

53. H. Brockman, Lipid monolayers: why use half a membrane to characterize protein-membrane interactions? Curr Opin Struct Biol 9, 438–443 (1999).

54. M. Dahim et al., Physical and photophysical characterization of a BODIPY phosphatidylcholine as a membrane probe. Biophys J 83, 1511–1524 (2002).

55. G. Gerebtzoff, A. Seelig, In silico prediction of blood-brain barrier permeation using the calculated molecular cross-sectional area as main parameter. J Chem Inf Model 46, 2638–2650 (2006).

56. A. Seelig, G. Gerebtzoff, Enhancement of drug absorption by noncharged detergents through membrane and P-glycoprotein binding. Expert Opin Drug Metab Toxicol 2, 733–752 (2006).

57. A. Witkowska, L. Jablonski, R. Jahn, A convenient protocol for generating giant unilamellar vesicles containing SNARE proteins using electroformation. Sci Rep 8, 9422 (2018).

58. W. Rawicz, K. C. Olbrich, T. McIntosh, D. Needham, E. Evans, Effect of chain length and unsaturation on elasticity of lipid bilayers. Biophys J 79, 328–339 (2000).

59. W. Rawicz, B. A. Smith, T. J. McIntosh, S. A. Simon, E. Evans, Elasticity, strength, and water permeability of bilayers that contain raft microdomain-forming lipids. Biophys J 94, 4725–4736 (2008).

60. R. Kwok, E. Evans, Thermoelasticity of large lecithin bilayer vesicles. Biophys J 35, 637–652 (1981).

61. T. Baumgart et al., Large-scale fluid/fluid phase separation of proteins and lipids in giant plasma membrane vesicles. Proc Natl Acad Sci U S A 104, 3165–3170 (2007).

62. T. M. Suchyna et al., Identification of a peptide toxin from Grammostola spatulata spider venom that blocks cation-selective stretch-activated channels. J.Gen.Physiol 115, 583–598 (2000).

63. T. M. Fischer, Bending stiffness of lipid bilayers. I. Bilayer couple or single-layer bending? Biophys J 63, 1328–1335 (1992).

64. T. Starke-Peterkovic et al., Cholesterol effect on the dipole potential of lipid membranes. Biophys J 90, 4060–4070 (2006).

65. H. F. Hofbauer et al., The molecular recognition of phosphatidic acid by an amphipathic helix in Opi1. J Cell Biol 217, 3109–3126 (2018).

66. E. E. Kooijman et al., What makes the bioactive lipids phosphatidic acid and lysophosphatidic acid so special? Biochemistry 44, 17007–17015 (2005).

67. A. Zemel, A. Ben-Shaul, S. May, Modulation of the spontaneous curvature and bending rigidity of lipid membranes by interfacially adsorbed amphipathic peptides. J Phys Chem B 112, 6988–6996 (2008).

68. D. Koller, K. Lohner, The role of spontaneous lipid curvature in the interaction of interfacially active peptides with membranes. Biochim Biophys Acta 1838, 2250–2259 (2014).

69. A. J. Sodt, R. W. Pastor, Molecular modeling of lipid membrane curvature induction by a peptide: more than simply shape. Biophys J 106, 1958–1969 (2014).

70. X. Yang et al., Structure deformation and curvature sensing of PIEZO1 in lipid membranes. Nature 604, 377–383 (2022).

71. C. S. Chiang, A. Anishkin, S. Sukharev, Gating of the large mechanosensitive channel in situ: estimation of the spatial scale of the transition from channel population responses. Biophys.J. 86, 2846–2861 (2004).

72. K. Vanommeslaeghe et al., CHARMM general force field: A force field for drug-like molecules compatible with the CHARMM all-atom additive biological force fields. J Comput Chem 31, 671–690 (2010).

73. L. Kalé et al., NAMD2: Greater Scalability for Parallel Molecular Dynamics. Journal of Computational Physics 151, 283–312 (1999).

74. A. D. MacKerell, Jr., M. Feig, C. L. Brooks, 3rd, Improved treatment of the protein backbone in empirical force fields. J Am Chem Soc 126, 698–699 (2004).

75. T. Darden, D. York, L. Pedersen, Particle Mesh Ewald - An N.Log(N) Method for Ewald Sums in Large Systems. Journal of Chemical Physics 98, 10089–10092 (1993).

76. M. Amblard, J. A. Fehrentz, J. Martinez, G. Subra, Methods and protocols of modern solid phase Peptide synthesis. Mol Biotechnol 33, 239–254 (2006).

77. Y. Zhao, M. C. Pirrung, J. Liao, A fluorescent amino acid probe to monitor efficiency of peptide conjugation to glass surfaces for high density microarrays. Mol Biosyst 8, 879–887 (2012).

## References

1. K. Vanommeslaeghe et al., CHARMM general force field: A force field for drug-like molecules compatible with the CHARMM all-atom additive biological force fields. J Comput Chem 31, 671–690 (2010).

2. L. Kalé et al., NAMD2: Greater Scalability for Parallel Molecular Dynamics. Journal of Computational Physics 151, 283–312 (1999).

3. A. D. MacKerell, Jr., M. Feig, C. L. Brooks, 3rd, Improved treatment of the protein backbone in empirical force fields. J Am Chem Soc 126, 698–699 (2004).

4. T. Darden, D. York, L. Pedersen, Particle Mesh Ewald - An N.Log(N) Method for Ewald Sums in Large Systems. Journal of Chemical Physics 98, 10089–10092 (1993).

5. W. Humphrey, A. Dalke, K. Schulten, VMD: visual molecular dynamics. J.Mol.Graph. 14, 33–38 (1996).

6. M. Amblard, J. A. Fehrentz, J. Martinez, G. Subra, Methods and protocols of modern solid phase Peptide synthesis. Mol Biotechnol 33, 239–254 (2006).

7. Y. Zhao, M. C. Pirrung, J. Liao, A fluorescent amino acid probe to monitor efficiency of peptide conjugation to glass surfaces for high density microarrays. Mol Biosyst 8, 879–887 (2012).

8. R. Gnanasambandam et al., GsMTx4: Mechanism of Inhibiting Mechanosensitive Ion Channels. Biophys J 112, 31–45 (2017).

9. W. Rawicz, B. A. Smith, T. J. McIntosh, S. A. Simon, E. Evans, Elasticity, strength, and water permeability of bilayers that contain raft microdomain-forming lipids. Biophys J 94, 4725–4736 (2008).

